# Subthreshold perturbation of DNA replication induces a secretory response and a bystander effect in naïve human fibroblasts

**DOI:** 10.64898/2026.08.20.745990

**Authors:** Benedetta Perdichizzi, Floriana Cappiello, Flavia Di Feo, Loredana Le Pera, Alfredo Pagliuca, Pasquale Valenzisi, Marco Rosina, Daniela Merlo, Annapaola Franchitto, Pietro Pichierri

## Abstract

Replication stress is a hallmark of cancer, where it drives DNA damage and genome instability. Yet subtle, subthreshold perturbations of DNA synthesis likely occur routinely in normal proliferating tissues, and their consequences for cell homeostasis are unknown. Using primary human fibroblasts, we show that doses of the DNA polymerase inhibitor aphidicolin, too low to engage the replication checkpoint, or produce detectable DNA breaks, nonetheless elicit low-level, ATM-dependent γH2AX phosphorylation uncoupled from overt damage. This near-silent perturbation reprograms gene expression, inducing replication-associated genes together with a discrete secretory programme dominated by matrix-remodelling proteases and matricellular factors. This output is not a senescence-associated secretory phenotype: the NF-κB/IL-1/IL-6 axis is co-ordinately repressed rather than induced, p53 target genes including CDKN1A are unchanged, and cells remain proliferative and non-senescent. Conditioned medium from exposed cells reproduces ATM-γH2AX activation in naïve fibroblasts without DNA damage, defining a “perturbed-replication bystander effect” (PeRBE). PeRBE is ROS-independent and mediated by heat-labile, proteinaceous factors, and in recipient cells it induces an extracellular-matrix programme that culminates in increased collagen production, without loss of proliferative capacity. A perturbation invisible to every standard replication-stress assay therefore generates a transmissible, protein-borne signal that instructs fibrogenic matrix remodelling in cells that never experienced it.

## INTRODUCTION

DNA replication can be perturbed by multiple events including competition with transcription, formation of secondary DNA structures or DNA repair intermediates (1, 2). Experimentally, replication stress (RS) - conventionally defined as persistent fork slowing, arrest or collapse - has most often been modelled by treatments with chemotherapeutic drugs, such as the topoisomerase I or II poisons camptothecin (CPT) and etoposide (ETP), or hydroxyurea (HU), an inhibitor of ribonucleotide reductase that depletes the cellular dNTP pool (3, 4). Using these set-ups, many steps of the RS response have been identified and mechanistically described, defining the main pathways that sense, arrest and process stalled forks (5–7). Nonetheless, the RS response has been essentially investigated and described as a fork-centred problem that cells manage internally through the checkpoint. Its classical outputs are cell-autonomous: cell cycle exit, mutations or genome instability and, eventually, cell death (1, 8–10).

More recently, consequences of RS that are not exclusively cell-autonomous have begun to be reported, including contributions to immune signalling through the cGAS-STING axis, NF-κB- mediated inflammation and senescence (11–14). However, all these consequences involve a substantial perturbation of DNA replication and the formation of detectable DNA damage, which acts as the real “second messenger” of the RS response beyond the fork.

In contrast, small perturbations of DNA replication - arising from subtle alterations in metabolism, from reduced fitness of replication caretakers carrying hypomorphic variants, or from the imbalance of proliferation genes that characterises very early precancerous lesions - would be undetectable by standard assays, yet be present and physiologically relevant. Cells appear able to detect a subtle arrest of replication forks and to mount a response even in the absence of the canonical checkpoint (15, 16), suggesting that the underlying molecular mechanisms operate locally without a threshold, but behave globally as a rheostat: a single local event does not impact cell homeostasis unless a global threshold is attained. Signalling and enforcement can therefore be uncoupled, and a perturbation may be sensed without any downstream execution being engaged. Whether a perturbation below this enforcement threshold has any non-cell-autonomous consequence has not, to our knowledge, been asked.

Instead, what is known so far are non-cell-autonomous consequences driven by paracrine phenotypes associated with extensive DNA damage or with senescence: the radiation-induced bystander effect (RIBE), which is often recapitulated by treatment with chemotherapeutics such as etoposide, and the senescence-associated secretory phenotype (SASP), which is also largely recapitulated by oncogene induction (17–21). RIBE is driven by DNA damage and involves production of ROS, direct cell-to-cell contact through gap junctions and secretion of inflammatory interleukins (18, 19, 22); its bystander outputs are notably the formation of DNA damage and, eventually, cell death (18). SASP is linked to senescence and p53/p21 signalling, and involves chronic activation of the DNA damage response, expression of the NF-κB-IL-1/IL-6 core and secretion of a bulk of interleukins that mediate bystander effects driving inflammation and fibrosis (21, 23). Both phenomena are therefore defined by the same prerequisite: damage severe enough to arrest or to kill the cell that transmits the signal. On this basis, only genuine RS, by elevating DNA damage, would be expected to elicit an inflammatory, SASP-like secretory response capable of affecting naïve cells and of driving pathological states such senescence, inflammation and also fibrosis in some instances. Fibrosis is linked to fibroblast activation in tissues (24, 25). Fibroblasts are the cell type populating the stroma and are responsible for extracellular matrix (ECM) metabolism (26, 27). Their activation leads to upregulation of several genes encoding ECM components, including collagens, and while controlled ECM production is part of tissue repair, its uncontrolled generation is associated with fibrosis in several pathological settings (24, 26, 28, 29).

Here, we investigated whether a subthreshold perturbation of DNA replication, induced globally using a very low dose of the polymerase inhibitor aphidicolin, below the conventional common fragile sites (CFS)-inducing range, could affect fibroblast homeostasis. Surprisingly, we found that such a perturbation - undetected by the checkpoint and producing only a minimal delay of fork progression - is nonetheless sufficient to modulate the expression of hundreds of genes and to establish a secretory phenotype distinct from SASP. This secretory phenotype exerts a bystander effect, which we called PeRBE (<u>Pe</u>rturbed <u>R</u>eplication <u>B</u>ystander <u>E</u>ffect), transcriptionally reprogramming naïve fibroblasts towards ECM production. Comparison with a genuine DNA damage response elicited by etoposide showed that the two states are transcriptionally opposite, establishing PeRBE as a distinct output of the perturbed replication response rather than a milder version of the damage response.

## RESULTS

### Minimal replication interference elicits a low-level ATM-H2AX activation without checkpoint engagement and DNA damage

Genuine replication stress (RS) is known to activate inflammatory and immune-related pathways (30, 31), and high doses of the radiomimetic drug Etoposide can induce a bystander effect (32). Before analysing the consequences of a subthreshold perturbation of DNA replication, we first assessed whether such mild interference engages canonical RS responses or induces DNA damage. Indeed, perturbing replication with high doses of HU or aphidicolin (APH), as well as with CPT or ETP, results in fork stalling or collapse, accumulation of ssDNA, checkpoint activation, DNA damage formation, and cell-cycle exit – either transiently or permanently – through quiescence or senescence (33) that are phenotypes that can trigger paracrine responses.

To avoid confounding effects from spontaneous senescence, we used a strain of human foreskin primary fibroblasts at early passages. These cells display a maximum of 5% SA-β-Gal positivity when used up to passage 28 (PD = 34), with more than 80% of cells actively proliferating at this stage (Figure S1A, B). For all experiments, cells were used between passages 16 and 23, when they can be defined as “young.” Treatment with 50 or 100 nM APH – doses that are far below those inducing CFS expression – for 24 h resulted in only a very mild reduction in replication fork speed, as assessed by DNA fiber assays (Figures 1A and S2A). The 50 nM dose did not induce detectable changes in global fork progression, whereas 100 nM APH caused approximately 10% slowing relative to untreated cells (Figures 1A-B). As a comparison, treatment with 400 nM APH - a low dose but sufficient to induce expression of CFS - reduced fork speed by about half (Figure 1A, B and S2A). Consistent with these effects, 50 nM APH did not alter the fraction of asymmetric forks, while 100 nM APH increased fork asymmetry by ∼1.5-fold, and 400 nM APH more than doubled the number of asymmetric forks relative to untreated cells (Figure 1C). Overall replication output remained comparable to control under subthreshold APH treatment, as shown by EdU incorporation at the end of the treatment while genuine RS induced by treatment with CPT resulted in an expected decrease in replicating cells (Figure S2B).

**Figure 1.**
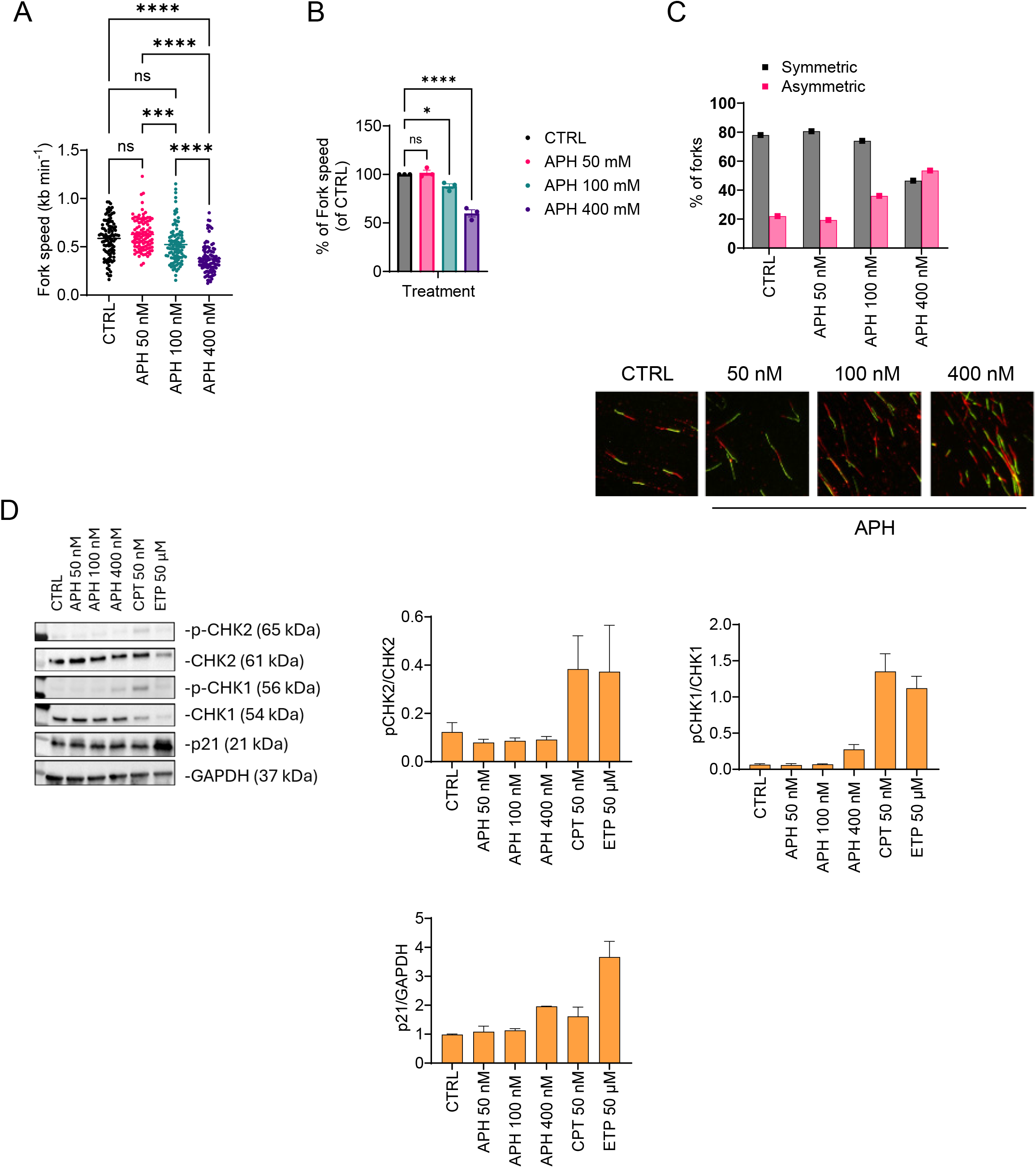
Subthreshold APH mildly perturbs replication fork dynamics without engaging the checkpoint. **(A)** Replication fork speed measured by DNA fiber assay in cells treated with 50, 100, or 400 nM APH for 24 h, or untreated (CTRL). Dual labelling with CldU and IdU – 30min each – was given at the end of treatment. Each dot represents a single fiber, pooled from n ≥ 3 independent experiments. Statistical significance was determined by one-way ANOVA. **(B)** Fork speed normalized to CTRL (%) under the same conditions, shown as mean ± SEM from n ≥ 3 independent experiments. Statistical significance was determined by one-way ANOVA. **(C)** Fraction of symmetric vs asymmetric forks under the same treatments, from n ≥ 3 independent experiments, with representative DNA fiber images (CldU/IdU) shown below. **(D)** Western blot analysis of p-CHK2 (T68), CHK2, p-CHK1 (S345), CHK1, and p21 (GAPDH as loading control) in untreated fibroblasts (CTRL) or cells treated with APH (50, 100, 400 nM), CPT (50 nM), or ETP (50 µM) for 24 h. Densitometric quantification of p-CHK2/CHK2, p-CHK1/CHK1, and p21/GAPDH ratios from n ≥ 3 independent experiments is shown; p21 levels were expressed relative to untreated control. In all panels, ns = not significant (p > 0.05); * p < 0.05; ** p < 0.01; *** p < 0.001; **** p < 0.0001.

We next asked whether this subthreshold perturbation of replication was detected by the replication checkpoint. To this aim, we evaluated phosphorylation of the checkpoint effector kinases CHK1 and CHK2, as well as activation of the p53-dependent axis by assessing p21 levels by Western blot. CPT and ETP served as positive controls. As shown in Figure 1D, subthreshold doses of APH failed to induce phosphorylation of CHK1 or CHK2 above basal levels, whereas CPT and ETP significantly enhanced phosphorylation of both kinases. Notably, mild replication stress associated with CFS expression, as induced by APH in the 400 nM range, was sufficient to induce low-level CHK1 phosphorylation, as previously reported (34). Consistent with activation of checkpoint effectors, p21 levels were significantly increased only in ETP-treated cells.

Checkpoint responses at perturbed replication forks are associated with formation of ssDNA (35). When we analysed exposure of parental ssDNA – an intermediate forming at stalled forks or at DNA gaps behind the fork (36, 37) – we detected only background levels in cells treated with subthreshold APH, consistent with the absence of checkpoint activation (Figure S3).

We next sought to confirm that subthreshold APH does not induce a DNA damage response (DDR) after 24 h. Immunofluorescence for pS1981-ATM, γH2AX, and pS824-KAP1 revealed that 100 nM APH significantly increased ATM and H2AX phosphorylation, whereas pKAP1 levels remained unchanged (Figure 2A). As expected, CPT and ETP induced more substantial phosphorylation of ATM, H2AX, and KAP1. Importantly, the low-level ATM-H2AX activation observed with 100 nM APH was unrelated to DNA damage, as both neutral and alkaline Comet assays failed to detect breaks under these conditions, while CPT produced readily detectable DNA damage (Figure 2B). Because low doses of APH are known CFS inducers, we also evaluated chromosomal damage in metaphase spreads from cells treated for 24 h with 100 nM or 400 nM APH. As expected, 400 nM APH resulted in chromosome breaks – most likely occurring at CFS as previously shown (ref PMID: 32966795) – whereas treatment with 50 or 100 nM APH, a four-fold lower dose, did not induce any detectable chromosomal lesions (Figure 2C).

**Figure 2.**
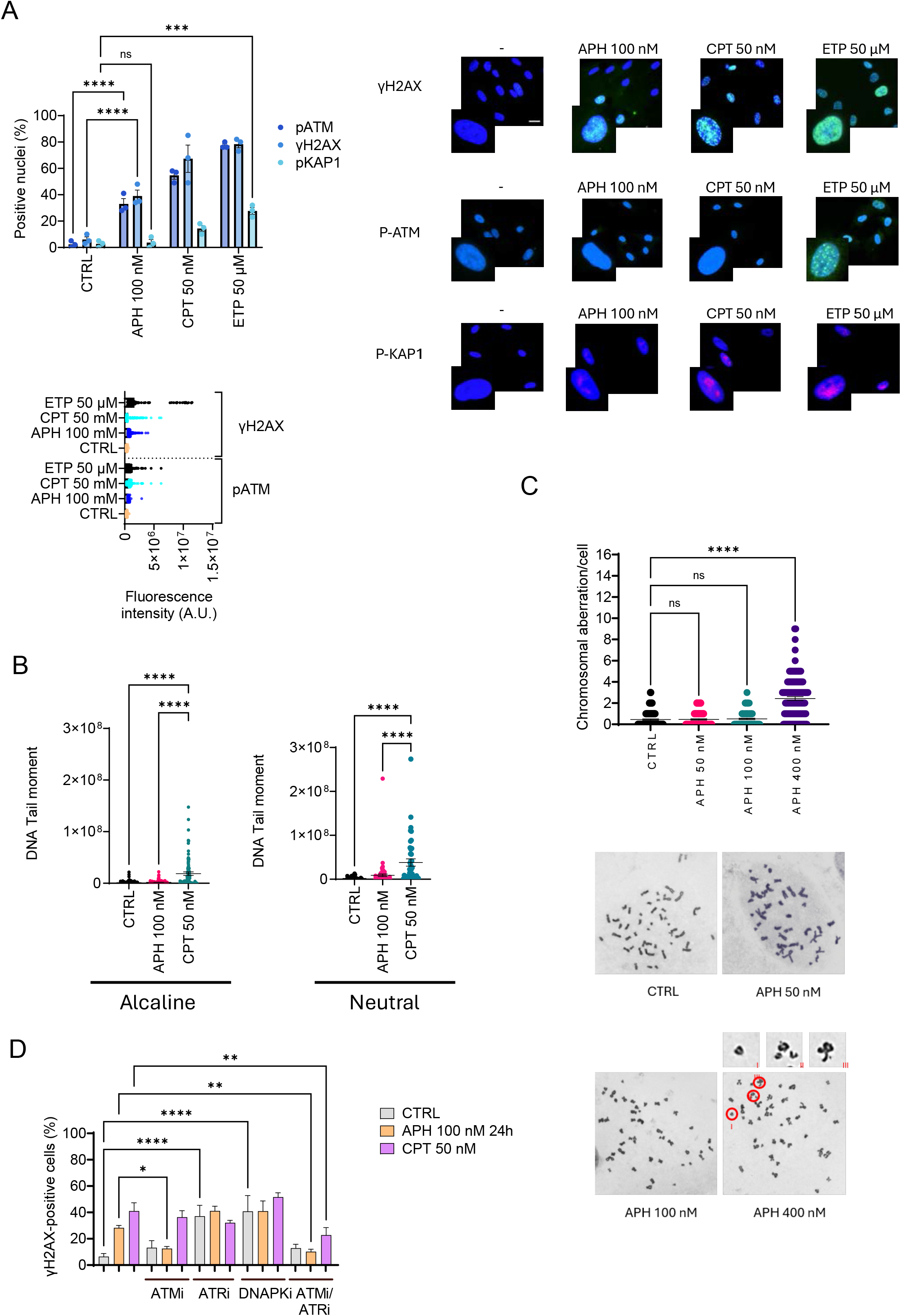
Subthreshold perturbed replication triggers ATM-dependent H2AX phosphorylation without inducing DNA damage. **(A)** Quantification of pS1981-ATM, γH2AX, and pS824-KAP1 positive nuclei (%) in cells treated with 100 nM APH, 50 nM CPT, or 50 µM ETP for 24 h (top), and corresponding quantification of fluorescence intensity (A.U.) for the same markers (bottom), with representative immunofluorescence images shown on the right. **(B)** DNA damage assessed by alkaline and neutral Comet assays (tail moment) following treatment with 100 nM APH or 50 nM CPT. **(C)** Chromosomal aberrations quantified in metaphase spreads from cells treated with 50, 100, or 400 nM APH for 24 hours. Representative images are shown (arrows indicate aberrations). **(D)** Quantification of γH2AX- positive cells (%) after treatment with 100 nM APH in the presence of ATM (ATMi, 10 µM), ATR (ATRi, 10 µM), and/or DNA-PK (DNA-PKi, 1 µM) inhibitors, added alone or in combination concurrently with the treatment. Data are representative of at least three independent experiments. Statistical significance was determined by one-way ANOVA; ns, not significant; * p < 0.05, ** p < 0.01; *** p < 0.001; **** p < 0.0001.

Our data therefore indicate that subthreshold perturbation of DNA replication is sufficient to trigger phosphorylation of ATM and H2AX in the absence of detectable DNA damage. To test whether H2AX phosphorylation under these conditions was ATM-dependent, we performed γH2AX immunostaining in cells treated with 100 nM APH in the presence or absence of ATMi, DNA-PKi, ATRi, or combinations thereof. Inhibition of ATM reduced by approximately half the fraction of γH2AX-positive cells following APH treatment, whereas DNA-PK inhibition had no appreciable effect. As expected, ATR inhibition caused a paradoxical increase in γH2AX-positive cells (Figure 2D). Notably, combined inhibition of ATM and ATR – but not of ATM and DNA-PK – abolished the H2AX phosphorylation induced by subthreshold replication stress (Figure 2D).

Collectively, these results indicate that a subthreshold perturbation of DNA replication, despite failing to induce DNA damage or checkpoint activation, is sufficient to trigger low-level ATM-dependent H2AX phosphorylation.

### Subthreshold perturbation of DNA replication induces a selective secretory program without senescence or p53 activation

Having shown that a subthreshold perturbation of DNA replication activates ATM–H2AX in the absence of other canonical checkpoint activation, detectable DNA damage or cell-cycle arrest, we asked whether it also alters cell homeostasis more broadly. Secretory responses are well documented downstream of DNA damage, oncogene-induced replication stress and senescence (20, 38, 39), but whether a perturbation below the threshold for checkpoint enforcement is sufficient to reprogram the secretome has not been addressed. We therefore performed RNA sequencing on primary human fibroblasts proliferating in the absence of treatment or exposed to 100 nM APH for 24 h (n = 4 independent biological replicates per condition; Figure 3A). Principal component analysis separated the two conditions along PC1, which accounted for 61.4% of the total variance, with no outlying sample and more than 13000 genes tested (Figure S4). Of the genes passing expression filtering, 2014 were differentially expressed (Benjamini–Hochberg FDR < 0.05, |log₂FC| > 0.585), comprising 1,024 up-regulated and 990 down-regulated genes; applying a more stringent |log₂FC| ≥ 1 threshold (at least a two-fold change) returned 638 up-regulated and 394 down-regulated genes (Figure 3B and Supplementary Table 1). The volcano plot distribution (Figure 3B) shows that, while the stringent cut-off isolates a core of high-magnitude changes, the statistically significant gene population spans a considerably broader response. Gene Ontology (GO) (40) over-representation analyses performed on protein-coding DEGs revealed that the up-regulated list was dominated by chromosome segregation (5.2-fold), DNA replication (4.7-fold) and cell division, and the down-regulated list by steroid and lipid biosynthesis and by extracellular structure organisation (Figure S5).

**Figure 3.**
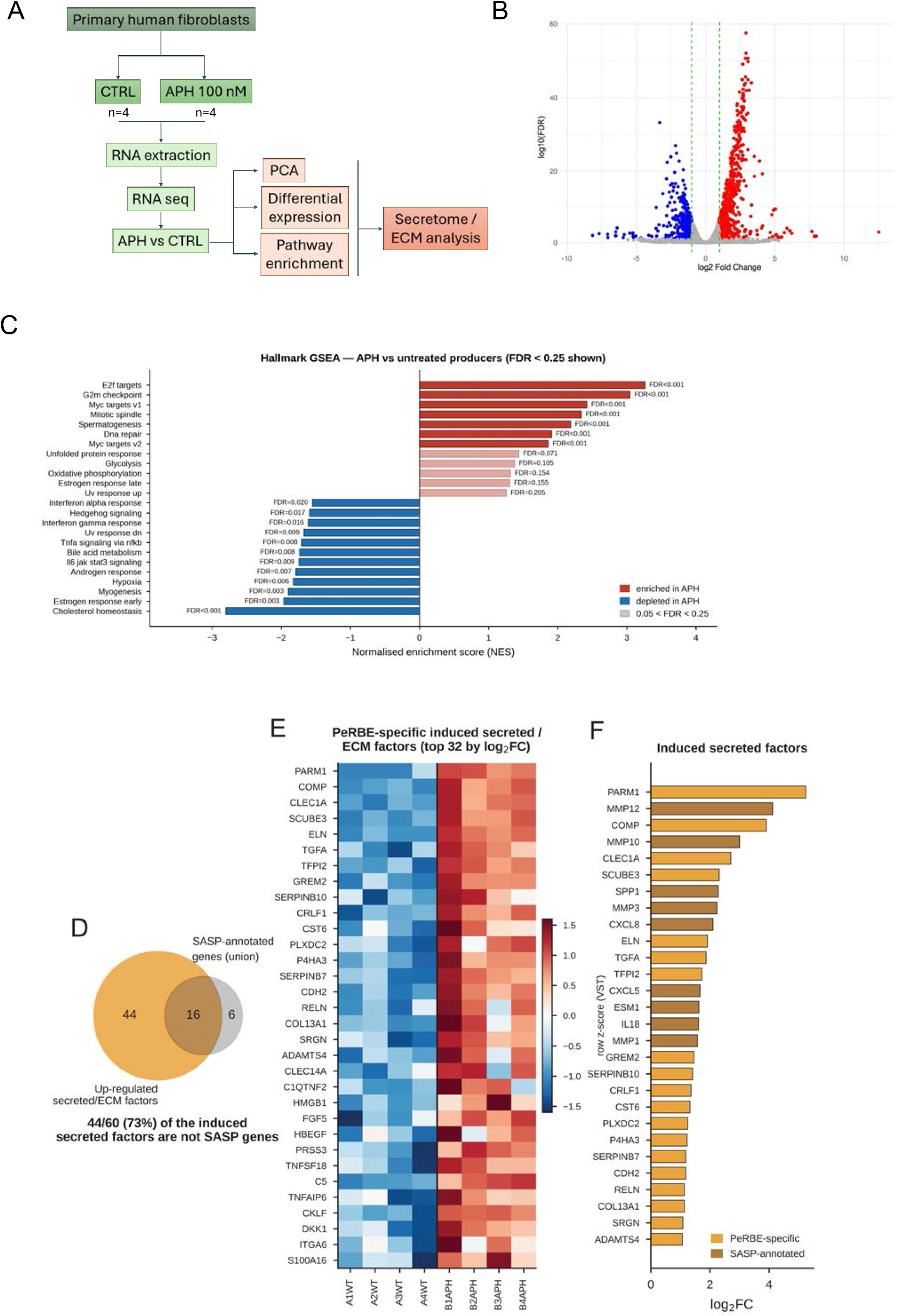
Subthreshold replication stress reprogrammes the transcriptome and induces a discrete secretory programme. **(A)** Experimental workflow: primary human fibroblasts were left untreated (CTRL) or treated with 100 nM APH for 24 h (n = 4 independent biological replicates per condition), followed by RNA extraction and RNA sequencing. Differential expression, pathway enrichment, and secretome/ECM analyses were performed on the APH vs CTRL comparison. **(B)** Volcano plot of differentially expressed genes (APH vs CTRL; FDR < 0.05, |log₂FC| > 2); up-regulated genes in red, down- regulated in blue. **(C)** Hallmark gene set enrichment analysis (GSEA) comparing APH-treated to untreated producer cells (FDR < 0.25 shown). Bars represent normalised enrichment score (NES); red, enriched in APH; blue, depleted in APH. **(D)** Pie chart showing that 44 of 60 (73%) up-regulated secreted/ECM factors are not annotated as SASP genes. **(E)** Heatmap of the top 32 PeRBE-specific induced secreted/ECM factors, ranked by Log₂FC, across CTRL and APH biological replicates. **(F)** Bar chart of induced secreted factors ranked by Log₂FC, classified as PeRBE-specific or SASP- annotated.

Gene set enrichment analysis confirmed that the most strongly enriched programmes in APH-treated cells were proliferative and replication-associated: E2F targets (NES 3.27, FDR < 0.001), G2M checkpoint (NES 3.05), MYC targets (NES 2.43) and DNA repair (NES 1.91) (Figure 3C). Of a curated 198-gene replication-stress panel, 101 genes were up-regulated and 2 down-regulated (odds ratio 13.9, P = 2 × 10⁻⁶⁰), including the Fanconi anaemia pathway, fork-associated factors and dNTP supply (Figure S5C and Supplementary Table 1). In contrast, neither the apical checkpoint machinery and DNA damage sensors nor the fork-remodelling factors were transcriptionally regulated, indicating that a response at the level of fork transactions or damage sensing is enacted post-translationally (Figure S5). Canonical proliferation genes were induced (including E2F1, FOXM1), as were replication-stress and genome-maintenance genes, most of which belong to the proliferation programme under the control of the E2F1 axis. Consistently, transcription-factor motif analysis was led throughout by E2F-family motifs (NES 2.33, FDR < 0.001; Figure S6). A further co-ordinate change accompanied this response: 62 of 66 detected replication-dependent histone genes were induced and none repressed (median log₂FC +2.35; odds ratio 200, P = 5 × 10⁻⁶⁵), together with the induction of histone chaperones and chromatin writers as a class (NES 1.65; ASF1B +2.22, UHRF1 +1.42, SUV39H1/2 +1.22 as the top upregulated, and EZH2, DNMT3B, HAT1) (Figures S6A and S7). Because the fraction of S-phase cells in the APH-treated population was not significantly increased relative to control (Figure S1), this proliferative and replication-associated signature is not a secondary consequence of an increased S-phase fraction.

The subthreshold APH treatment also induced a discrete set of secreted and matrix-associated factors. To define this compartment, we intersected the differentially expressed genes with the Human Protein Atlas predicted secretome (2,520 genes, curated from UniProt subcellular localisation annotations combined with signal peptide and transmembrane predictions) and with genes annotated under the Gene Ontology term ’extracellular region’ (GO:0005576), which returned a prioritised subset of 118 candidate secreted factors. Functional enrichment of this subset showed a pronounced over- representation of matrix metalloproteases, ECM constituents and collagen catabolic and metabolic processes (Figure S8). Notably, this occurred against a background of global repression of matrix gene expression (NES −2.18, FDR < 0.001; 25% of matrisome genes down-regulated; Figure S5 and Supplementary Table 1). Of the 1,024 up-regulated genes, 60 encode secreted or extracellular-matrix proteins, dominated by matrix-remodelling proteases and their regulators (MMP12 +4.12, MMP10 +3.00, MMP3 +2.24, MMP1 +1.58, ADAMTS4 +1.07, TIMP1 +1.05, PLAU +0.84, PLAUR +0.67, SERPINE1 +0.61), together with structural and matricellular components (COMP +3.91, SCUBE3 +2.31, ELN +1.91, TNC +0.75, POSTN +0.60, MFAP5 +0.73, PLOD2 +0.61, LRRC15 +0.68), growth factors and morphogen modulators (TGFA +1.87, HBEGF +0.90, GREM2 +1.45, DKK1 +0.80, CRLF1 +1.36), and protease inhibitors and damage-associated factors (TFPI2 +1.73, CST6 +1.32, HMGB1 +0.98, ANXA1 +0.69, TNFAIP6 +0.81) (Figure 3D–F; Supplementary Table 2). At the transcription level, a subthreshold perturbation of DNA replication therefore establishes a discrete secretory phenotype superimposed on a globally repressed matrix programme.

To determine whether this secretory phenotype corresponds to a senescence-associated secretory phenotype (SASP), we resolved a curated core SASP gene set (75 genes detected) into functional modules (Figure 4). Induction was almost entirely confined to the protease and matrix-remodelling arm (MMP12, MMP10, MMP3, MMP1, TIMP1, PLAU, PLAUR, SERPINE1), together with three chemokines (CXCL8, CXCL5, CXCL2) and IL18, IL11 and SPP1. The NF-κB–IL1–IL6 axis that defines the SASP was not engaged: IL6 (−0.44), IL1B (+0.21), CXCL1 (+0.30) and CCL2 (+0.23) were unchanged and IL1A was not detectable. No TGF-β ligand was induced (TGFβ1 +0.04, TGFβ2 −0.33, both n.s.; TGFβ3 −0.73, FDR = 1.3 × 10⁻⁵). At the level of whole programs, the discordance was unambiguous: TNFα signalling via NF-κB (NES −1.70), IL6/JAK/STAT3 signalling (−1.75), inflammatory response (−1.32), IFN-γ (−1.61) and IFN-α (−1.55) responses were all significantly depleted in APH-treated cells (Figure 4). Consequently, a conventional enrichment test against the SASP gene set, as a whole, returns a significant overlap that is driven entirely by its protease arm and masks the co-ordinate repression of its inflammatory core. In contrast, the genuine RS and DNA damage induced by 24 h of 50 nM ETP produced a response dominated by inflammatory and immune-related genes (Figure S9 and Supplementary Table 3). Subthreshold replication perturbation therefore induces a secretory output that overlaps the SASP only in its protease arm while repressing the program that defines it.

**Figure 4.**
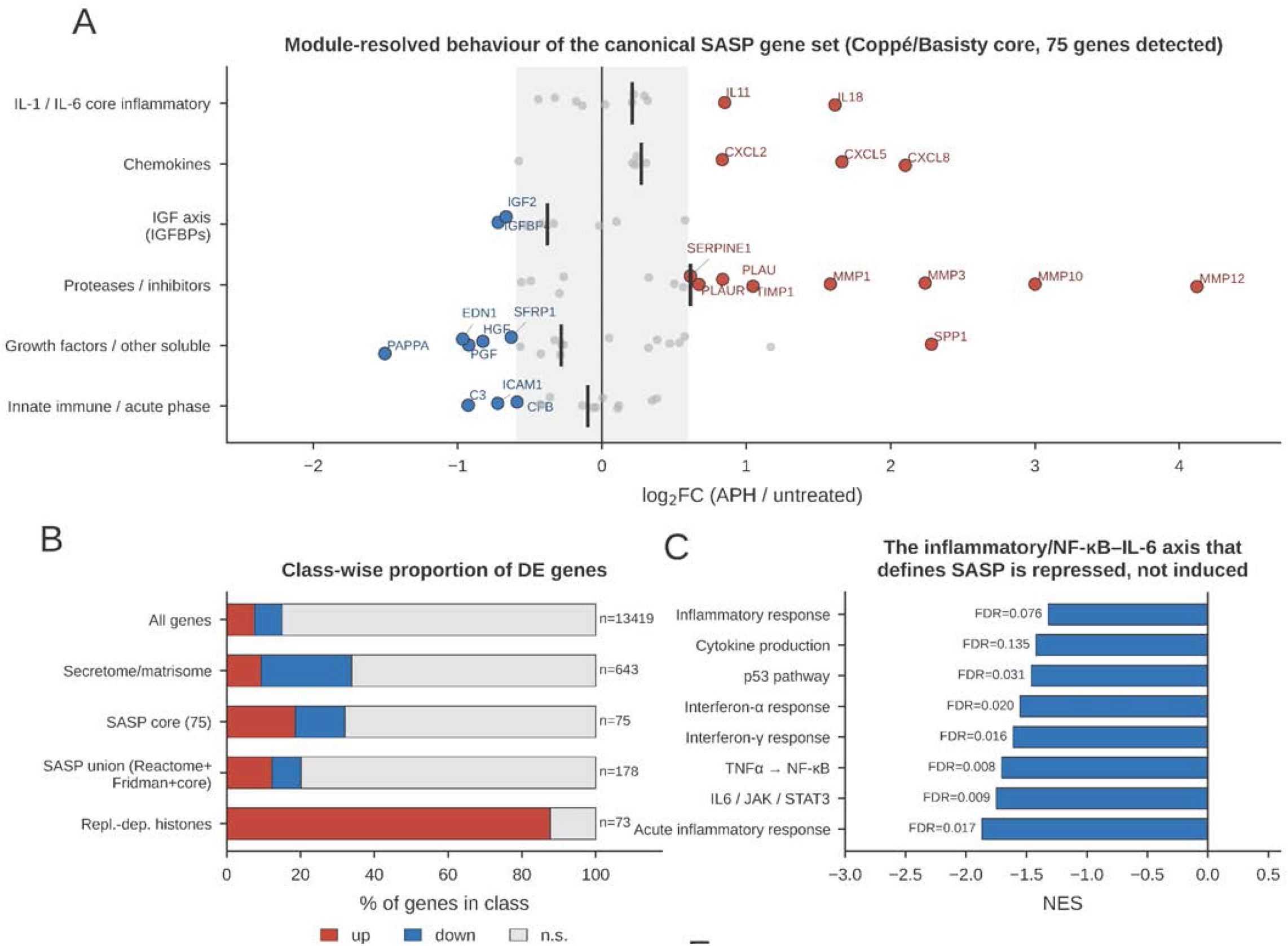
The induced secretory programme is dissociated from the canonical inflammatory SASP. **(A)** Module-resolved behaviour of the canonical core SASP gene set (Coppé/Basisty; 75 genes detected), grouped by functional category (IL-1/IL-6 core inflammatory, chemokines, IGF axis/IGFBPs, proteases/inhibitors, growth factors/other soluble, innate immune/acute phase). Each dot represents a gene plotted by log₂FC (APH/untreated). **(B)** Class-wise proportion of differentially expressed genes (up, down, or not significant-ns) across gene classes: all genes, secretome/matrisome, SASP core (75 genes), SASP union (Reactome + Fridman + core, n = 178), and replication-dependent histones. **(C)** Normalised enrichment scores (NES) for hallmark inflammatory and NF-κB/IL-6 axis gene sets (inflammatory response, cytokine production, p53 pathway, interferon-α response, interferon-γ response, TNFα signalling via NF-κB, IL6/JAK/STAT3 signalling, acute inflammatory response).

Consistent with the absence of an inflammatory SASP, the p53-p21 axis was not activated. CDKN1A was unchanged (log₂FC +0.10) despite being abundantly expressed at baseline (∼9,600 normalised counts), as were CDKN2A (−0.14), TP53 (+0.36), MDM2 (−0.13) and GADD45A (+0.16); CDKN2B was significantly repressed (−0.84). Of 32 canonical p53 target genes only TP53I3 was induced, and the p53 program as a whole was significantly depleted (NES −1.46, FDR = 0.031) (Figure 5A-C). Furthermore, LMNB1, whose loss is a hallmark of senescence, was increased (+1.54), as was MKI67 (+2.07), indicating that producer cells retain proliferative identity. We tested this prediction functionally by assessing senescence by SA-β-Gal staining and EdU incorporation after 24 h of 100 nM APH. APH-treated cells showed no proxy of premature senescence and were indistinguishable from untreated controls, whereas senescent fibroblasts of the same strain showed a marked increase in SA-β-Gal positivity and reduced proliferative potential, as indicated by decreased EdU incorporation (Figure 5D, E).

**Figure 5.**
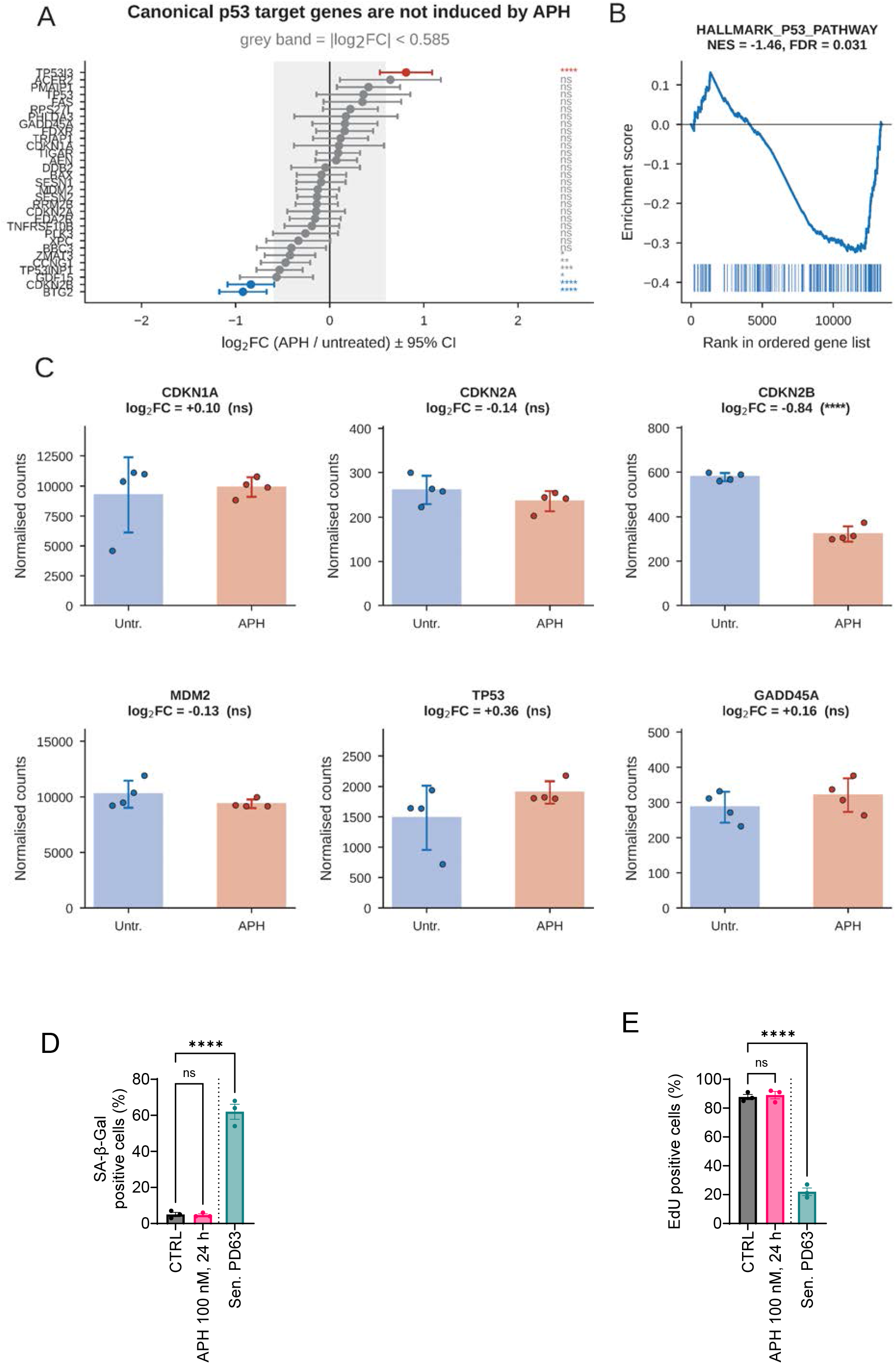
Subthreshold replication stress does not activate the p53–p21 axis or induce senescence. **(A)** Forest plot of canonical p53 target genes, showing log₂FC (APH/untreated) ± 95% CI; grey band indicates |log₂FC| < 0.585. **(B)** GSEA enrichment plot for the HALLMARK_P53_PATHWAY gene set (NES = −1.46, FDR = 0.031). **(C)** Normalised counts of representative p53-pathway genes (CDKN1A, CDKN2A, CDKN2B, MDM2, TP53, GADD45A) in untreated (Untr.) and APH-treated cells; Log₂FC and significance are indicated above each panel (ns, not significant; ****P < 0.0001). **(D)** Percentage of SA-β-Gal-positive cells in CTRL, APH-treated (100 nM, 24 h), and senescent (Sen. PD63) fibroblasts. **(E)** Percentage of EdU-positive cells in the same conditions as in (D), showing that APH-treated cells retain proliferative capacity, unlike senescent controls. Data are mean ± SD; ns, not significant; ****P < 0.0001.

Collectively, these results show that a perturbation of DNA replication below the threshold for checkpoint enforcement and in the absence of detectable DNA damage or cell-cycle arrest, is sufficient to reprogram the secretory output of primary human fibroblasts without engaging p53, without loss of proliferative identity and without the inflammatory program that defines the SASP. We refer to this condition hereafter as the perturbed-replication bystander effect (PeRBE) producer state.

### Subthreshold perturbation of DNA replication triggers a bystander ATM-H2AX activation

Cells experiencing a subthreshold perturbation of replication therefore acquire a secretory phenotype that is qualitatively distinct from those described downstream of overt DNA damage or senescence. We next asked whether this output is biologically active – that is, whether medium conditioned (CM) by these cells is sufficient to alter the behaviour of naïve fibroblasts. Since bystander effect from DNA damage involving activation of ATM can activate in trans ATM also in naïve cells, we first analysed phosphorylation status of S1981 of ATM and levels of γH2AX by immunofluorescence. Naïve young human fibroblasts of the same strain of the producer cells and matched for passage number, were exposed to CMs from control or APH-treated cells according to the protocol shown in Figure 6A (hereafter responder cells). As expected, pS1981-ATM immunofluorescence was clearly increased in producer cells treated with 100 nM APH, and increased further in cells treated with 50 nM CPT, which serves as a genuine RS and DNA damage control alongside ETP (Figures 6B and 2A). Notably, exposure to CM from APH-treated producers also induced an increase in the pS1981-ATM signal that was absent in cells exposed to control CM (Figure 6B). This bystander phosphorylation of ATM was confirmed by in situ proximity ligation assay between pS1981-ATM and ATM (Figure 6C). As shown in Figure 6C, the levels of phosphorylated H2AX were similarly increased by exposure to CM from APH-treated producers relative to control CM, and, consistent with ATM activation, H2AX phosphorylation was more evident in cells exposed to CM from CPT-treated producers (Figure 6B, C). Although the number of nuclei positive for ATM or H2AX phosphorylation was increased by both APH and CPT CM, the intensity of the immunofluorescence signal was increased only by APH CM. A comparable bystander H2AX phosphorylation following exposure to APH-conditioned media was confirmed in a second strain of young primary human fibroblasts (Ag21859 – passage number 12) (Figure S10A).

**Figure 6.**
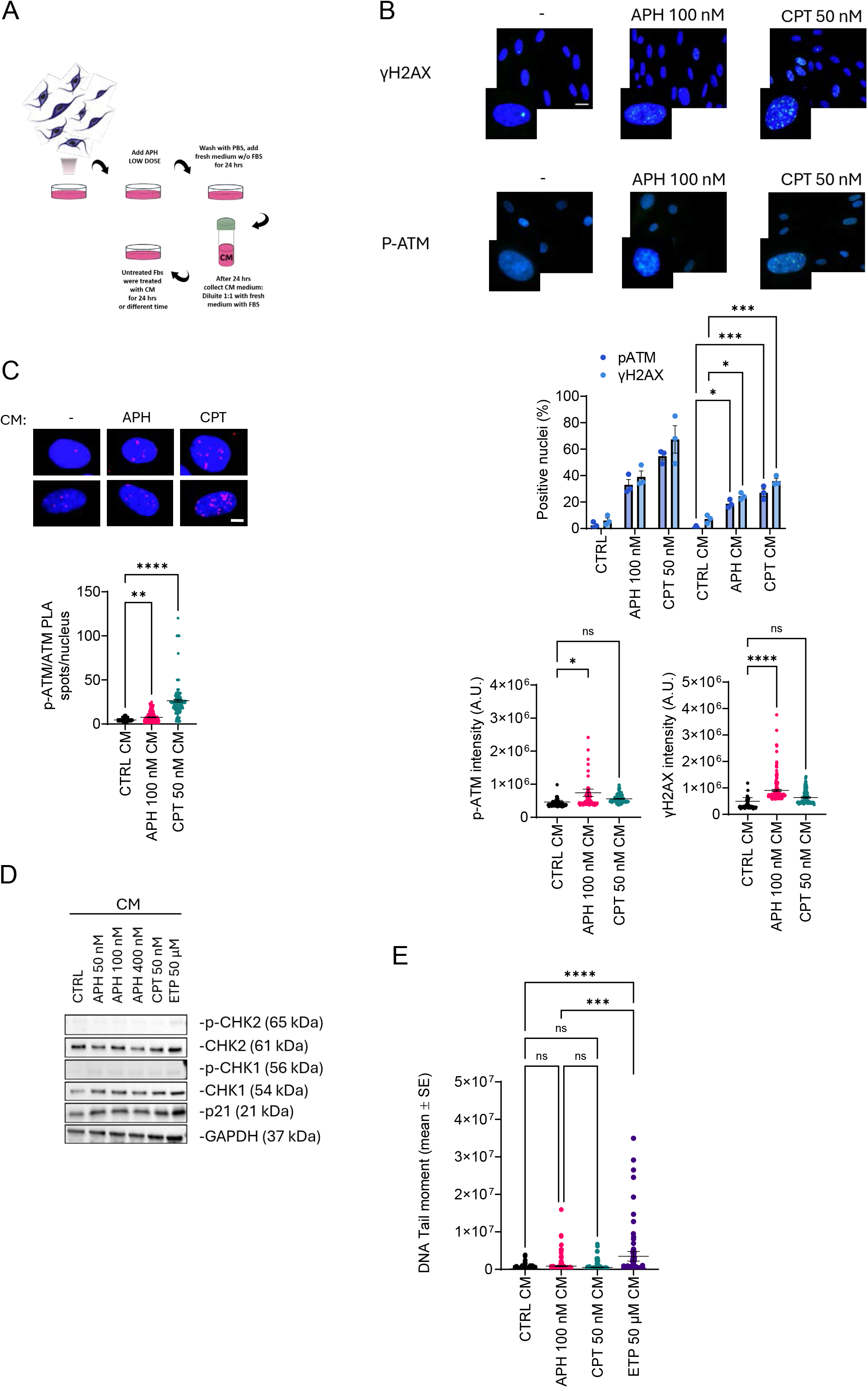
Subthreshold replication stress triggers bystander ATM–H2AX activation in naïve responder cells. **(A)** Experimental workflow: producer fibroblasts were treated with subthreshold dose APH, washed, and incubated in serum-free medium for 24 h; conditioned medium (CM) was then collected, diluted 1:1 with fresh serum-containing medium, and used to treat untreated (naïve) responder fibroblasts for 24 h before evaluating the endpoints. **(B)** Representative immunofluorescence images (top) and quantification (bottom) of γH2AX and p-ATM (S1981) in responder cells exposed to CM from untreated (CTRL), APH-treated (100 nM), or CPT-treated (50 nM) producer cells. Bar graphs show the percentage of positive nuclei and scatter plots show p-ATM and γH2AX signal intensity per nucleus. **(C)** Representative images (top) and quantification (bottom) of *in situ* PLA between mouse anti p-ATM (S1981) and rabbit anti-ATM in responder cells exposed to CM from control, APH-, or CPT-treated producers. Each dot represents the number of PLA spots per nucleus. **(D)** Western blot analysis of p-CHK2 (T68), CHK2, p-CHK1 (S345), CHK1, p21, and GAPDH (loading control) in responder cells exposed to CM from control or from producers treated with 50, 100, or 400 nM APH, 50 nM CPT, or 50 µM ETP for 24 h. **(E)** Quantification (left) and representative images (right) of DNA damage assessed by Neutral comet assay (tail moment) in responder cells exposed to CM from control, APH-, CPT-, or ETP-treated producers. In all experiments data are from 3 independent replicates and statistical significance is determined by one-way ANOVA: ns, not significant; * p < 0.05; **P < 0.01; ***P < 0.001; ****P < 0.0001.

To assess whether prolonging the subthreshold perturbation of replication in producers affects the bystander response, we monitored γH2AX levels by immunofluorescence in responder cells as a proxy for PeRBE. The fraction of γH2AX-positive responder cells decreased as the duration of APH treatment in producer cells increased (Figure S10B), indicating that the transmissible activity is transient rather than cumulative or suggesting that the prolonged subthreshold RS converts into a genuine RS/DNA damage changing the nature of the response as suggested for senescence-associated paracrine activity (41). The PeRBE-mediated phosphorylation of ATM and H2AX was not abrogated by treatment of the CM with the ROS scavenger NAC (Figure S11A), whereas mild heat inactivation of the CM at 65 °C substantially reduced the fraction of γH2AX-positive nuclei in responder cells (Figure S11B), indicating that the transmitted signal is ROS-independent and proteinaceous. Size fractionation of conditioned media from control, APH- and ETP-treated producers through 30 kDa and 100 kDa cut-off filters showed that biological activity - estimated by bystander γH2AX immunofluorescence - was retained in both fractions and was, if anything, increased in the ≤30 kDa fraction, suggesting that the mediators are predominantly low-molecular-weight (≤30 kDa) secreted proteins (Figure S11C).

To distinguish the bystander activation of ATM-H2AX in PeRBE from that induced by DNA damage, such as after ETP or ionising radiation, we assessed the activation of CHK1, CHK2 and p21 by Western blot in responder cells as markers of DNA damage and cell-cycle arrest. Comparing lysates from responder cells exposed to APH- or ETP-conditioned media, we found no phosphorylation of CHK1 or CHK2, and only a very modest increase in p21 restricted to cells exposed to ETP-derived CM (Figure 6D). Consistent with this, responder cells exposed to APH-derived CM did not show any DNA break by neutral or alkaline Comet assay, while exposure to ETP-derived CM produced DNA double-strand breaks (DSBs) in responder cells (Figure 6E).

Altogether, our results show that PeRBE is associated with bystanding activation of ATM-H2AX in naïve responder cells when they are exposed to CM, which is unrelated to DNA damage or cell cycle arrest. Furthermore, our results indicate that PeRBE is ROS-independent and transmitted by heat- label proteinaceous factors.

### PeRBE reprograms bystander cells towards an extracellular-matrix-producing phenotype without compromising their proliferation

Since PeRBE is able to induce in trans activation of ATM-H2AX in responder cells, we investigate whether the secretory phenotype associated with the producer arm of PeRBE was also able to induce any other functional output. To explore this possibility, we exposed responder cells for 24 h with CM conditioned by producers treated with 100 nM APH for 24 h, or by untreated producers, and profiled them by RNA sequencing (n = 4 per condition). Rather than assay a preselected endpoint, we used the profiling to nominate candidate functional outputs in an unbiased manner (Figure 7A). Gene set enrichment analysis showed that the programs induced in responder cells were overwhelmingly related to the extracellular matrix: extracellular matrix organization (NES 2.70, FDR < 0.001), collagen formation (2.54), collagen biosynthesis and modifying enzymes (2.56), ECM proteoglycans (2.62), the NABA matrisome (2.26) and a curated myofibroblast/fibrotic ECM signature (myCAF; 2.44), together with epithelial–mesenchymal transition (2.16), contractile/myofibroblast (2.11) and TGF-β signaling (2.05) (Figure 7B). Signature scores computed per replicate separated the two groups completely, with all four APH-CM samples exceeding all four controls for the myCAF/ECM, matrisome, collagen and TGF-β signatures (Figure 7C). One hundred and one matrix-annotated genes were significantly induced (FDR < 0.05, |log₂FC| > 0.585, mean normalised counts ≥ 100), including fibrillar and network collagens (COL1A1 +1.53, COL5A1 +1.34, COL15A1 +1.15, COL6A1 +1.10, COL1A2 +0.95, COL4A2 +0.91), proteoglycans and basement membrane components (HSPG2 +1.61, BGN +1.13, NID1 +1.07, NID2 +1.06, LAMA5 +1.06), matricellular and matrix-organizing proteins (TNXB +1.63, PRELP +1.39, AEBP1 +1.38, FBLN1 +1.10, TNC +1.04, MXRA5 +1.06) and latent TGF-β binding proteins (LTBP2 +1.22, LTBP3 +1.00, LTBP4 +0.98) (Figure 7D). A collagen-producing program therefore emerged as one of the dominant predicted outputs of the response, and we tested it directly.

**Figure 7.**
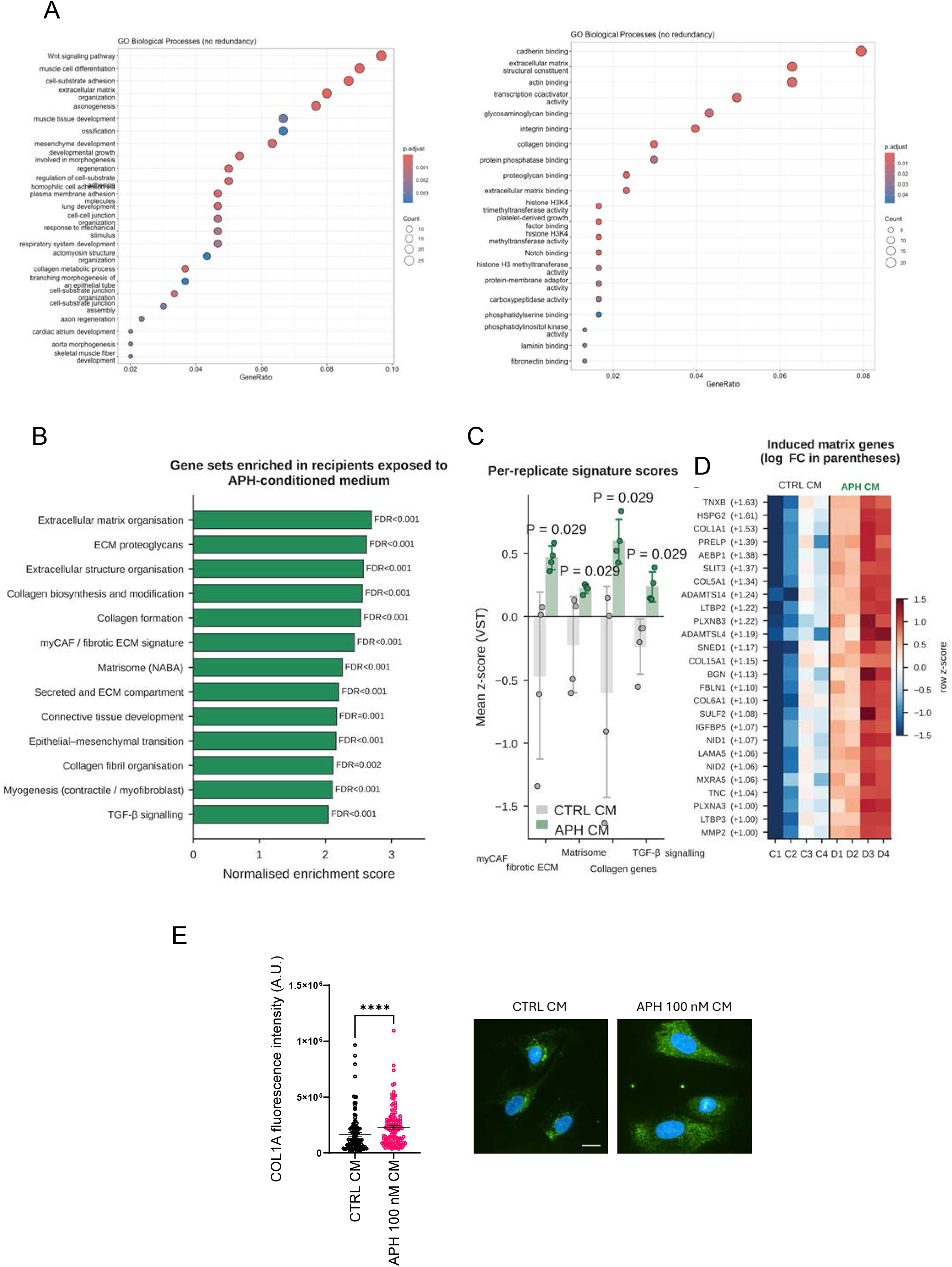
PeRBE-conditioned medium reprograms bystander fibroblasts towards an extracellular matrix/collagen gene expression program. RNA sequencing was performed on bystander fibroblasts exposed for 24 h to conditioned medium (CM) from APH-treated or untreated producer cells (n = 4 per condition). **(A)** GSEA dot plots showing the top non-redundant GO Biological Process (left) and Molecular Function (right) terms enriched in responders exposed to APH-CM. Dot size reflects gene ratio/count; colour indicates adjusted p-value. **(B)** Normalised enrichment scores for the top gene sets induced by APH-CM, including ECM organisation, collagen biosynthesis, the myCAF/fibrotic ECM signature, the NABA matrisome, EMT, and TGF-β signalling. FDR values are shown per bar. **(C)** Per-replicate signature scores (mean z-score, VST) for the myCAF/ECM, matrisome, collagen, and TGF-β signalling signatures in fibroblasts exposed to CTRL CM (grey) or APH CM (green). Each dot is a biological replicate (n = 4/group); bars show mean ± S.E.M. Mann–Whitney U test, P = 0.029. **(D)** Heatmap of the top 25 induced matrix genes (row z-score of normalised counts) across CTRL CM (C1–C4) and APH CM (D1–D4) replicates, ranked by fold change; Log₂FC shown in parentheses. **(E)** On the left, COL1A fluorescence was quantified by measuring the corrected total cell fluorescence (CTCF) for individual cells and expressed as COL1A fluorescence intensity in arbitrary units (A.U.). Data are presented as individual values with mean ± S.E.M. On the right, representative immunofluorescence images showing COL1A staining (green) in fibroblasts exposed to conditioned medium from untreated cells (CTRL CM) or from cells treated with aphidicolin (APH 100 nM CM) (scale bar: 20 μm). Nuclei were counterstained with DAPI (blue). Statistical significance was assessed using an unpaired two-tailed Student’s t-test; **** p < 0.0001.

Collagen production was evaluated by COL1A immunofluorescence comparing fibroblasts exposed for 48 h to CMs from APH-treated or untreated cells. As shown in Figure 7E, a limited COL1A staining was detectable in control fibroblasts. In sharp contrast and consistent with DGE analysis, anti-COL1A immunostaining was significantly increased in cells exposed to APH CM (Figure 7E). Myofibroblast and myCAF states are characteristically proliferative rather than arrested (29), and TGF-β signalling, which is among the induced PeRBE programs, is cytostatic in many fibroblast contexts (42). We therefore asked whether the matrix program in responder cells was accompanied by preserved replication competence, and whether it could be distinguished on that basis from a senescence-associated fibrotic state. To this end, we monitored the number of replication-competent cells in responder cells by EdU immunostaining and proliferation by Incucyte® SX5 live-imaging platform over time. A 30 min pulse with EdU followed by Click-It and evaluation of the number of S-phase cells, showed an absence of decline in the replication potential of responder cells at the end of exposure with CM from APH-treated cells (Figure 8A). If any, exposure to CM from APH-treated cells resulted in a mild but reproducible increase in the number of EdU-positive cells as compared with control CM (Figure 8A). Consistent with the small increase in the number of replicating cells, also proliferation over time was increased in bystander cells exposed to APH-conditioned media as compared to cells exposed to control CM (Figure 8B-D). This increased proliferation was apparent by the confluence area (proportional to the number of cells) and by the shorter time to inflection in the proliferation curve. Of note, the pro-cell growth effect of PeRBE is against any implication of deprivation of growth factors from the CM.

**Figure 8.**
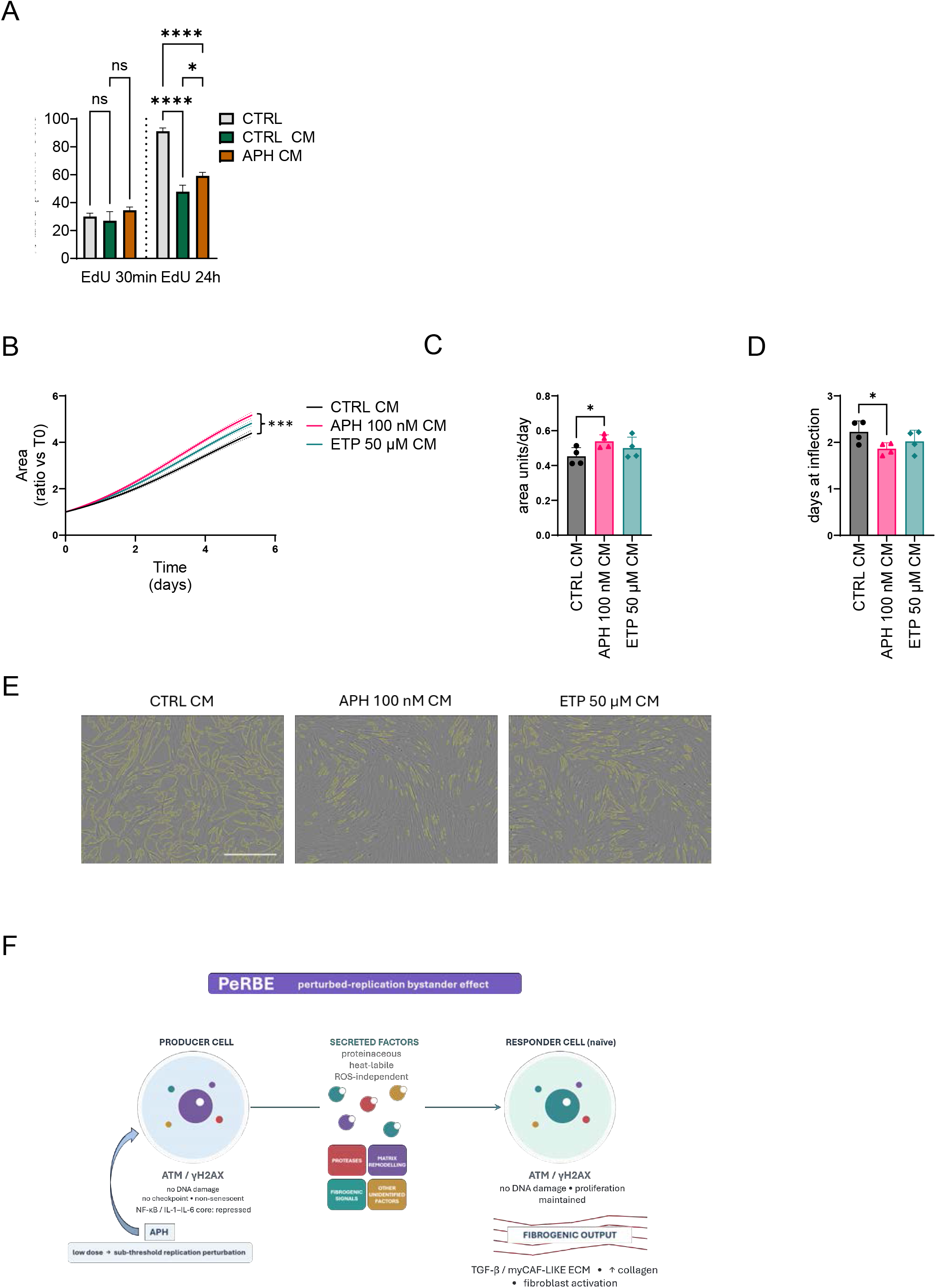
Conditioned media from APH-treated cells enhances bystander cell proliferation over time. **(A)** EdU incorporation in bystander fibroblasts, measured after a short 30 min pulse or continuous 24 h labelling at the end of CM exposure (CTRL vs. CTRL CM vs. APH CM). Data show mean ± S.E.M. One-way ANOVA; ****p < 0.0001, *p < 0.05, ns = not significant. **(B)** Cell proliferation kinetics over a 5-day period using the Incucyte system in bystander cells exposed to conditioned media (CM) derived from untreated control cells (CTRL CM), 100 nM Aphidicolin-treated cells (APH 100 nM CM), or 50 µM Etoposide-treated cells (ETP 50 µM CM). Data are expressed as cell area coverage relative to baseline (ratio vs T0). Statistical significance was determined by Two-Way ANOVA test (***p < 0.0001 APH 100 nM CM vs. CTRL CM at day 5). **(C)** Quantification of growth rate expressed as area units per day. Statistical significance was assessed using an unpaired two-tailed Student’s t-test (* p < 0.05 APH 100 nM CM vs. CTRL CM). **(D)** Flex point analysis showing the time (days) to inflection in the proliferation curves. Statistical significance was assessed using an unpaired two-tailed Student’s t-test (* p < 0.05 APH 100 nM CM vs. CTRL CM). **(E)** Representative phase-contrast live-cell images displaying cell density and coverage in bystander cells under the specified CM conditions at day 5. Scale bar = 400 µm. **(F)** Proposed model: A subthreshold perturbation of DNA replication in the producer cell (100 nM APH) triggers ATM-dependent γH2AX phosphorylation in the absence of checkpoint engagement or detectable DNA damage, while the NF- κB/IL-1/IL-6 axis is repressed, so producer cells remain non-senescent and proliferative. This state drives secretion of proteinaceous, heat-labile, ROS-independent factors (proteases, matrix-remodelling factors and fibrogenic signals) that activate ATM/γH2AX in naïve responder cells without DNA damage or loss of proliferation, culminating in a fibrogenic output marked by TGF-β activation, myCAF-like ECM remodelling and increased collagen production.

Taken together, these data show that the secretome generated by a subthreshold perturbation of DNA replication reprograms naïve fibroblasts towards an extracellular-matrix program associated with increased collagen expression, and that it does so in cells that remain fully replication-competent. The bystander arm of PeRBE therefore produces an ECM-producing phenotype uncoupled from growth arrest, distinguishing it from the fibrogenic states associated with senescence; the preservation of proliferation also argues against depletion of nutrients or growth factors from the conditioned medium as a trivial explanation. A perturbation of DNA replication that is too mild to engage the checkpoint or to damage the cell in which it occurs is thus sufficient to instruct a fibrogenic program in neighboring cells that have never experienced it.

## DISCUSSION

The response to replication stress has been studied almost exclusively as a cell-autonomous programme that preserves viability and genome integrity, and that requires a substantial perturbation or a block of DNA replication in order to be engaged and to deliver an output (1, 7). Here we show that a perturbation far below that threshold – a dose of aphidicolin that slows global fork progression by approximately 10%, does not induce CFS expression, does not activate the replication checkpoint and produces no detectable DNA damage – is nonetheless sufficient to generate a paracrine signal that instructs an ECM-producing phenotype in naïve, proliferating human primary fibroblasts that never experienced it. The perturbation is invisible to every assay conventionally used to define replication stress, yet it is neither silent in the cell in which it occurs nor confined to it.

The producer state defines what we would describe as signalling without enforcement of any classical RS response. Proliferating human fibroblasts respond to a minimal reduction in fork rate by activating ATM and phosphorylating H2AX, but this activation occurs without DNA damage, without the acknowledged proxies of RS such as ssDNA exposure, and without phosphorylation of the effector kinases CHK1 and CHK2. Downstream, the p53 program is not merely unengaged but significantly depleted (NES −1.46), CDKN1A is unchanged from the basal level, LMNB1 is increased, and the proliferative transcriptional programme is retained and indeed amplified through the E2F axis. The perturbation is therefore sensed and transduced, while none of the executive arms of the response is engaged. This dissociation has a consequence that reaches beyond the specific phenomenon we describe: if a subthreshold perturbation of replication produces a transcriptional and paracrine output while remaining negative for every standard cell-autonomous readout – no breaks, no arrest, no checkpoint markers – then the large body of “no effect” observations obtained at low doses may have been interrogating the wrong compartment. Thresholds, in this view, do not separate response from no response; they separate signalling from enforcement, and the sub-enforcement regime has outputs of its own.

How ATM is activated under these conditions remains open, and we regard it as the principal mechanistic question raised by this work. ATM can be activated independently of DSBs at reversed forks, at R-loops and in response to changes in chromatin topology (43–48). Two observations bear on this. First, ATM and H2AX phosphorylation occur without any detectable phosphorylation of KAP1, a canonical and highly sensitive ATM substrate at breaks (49), which argues against a cryptic population of DSBs below the detection limit of the Comet assay. Second, combined inhibition of ATM and ATR abolishes γH2AX, indicating that the residual signal is ATR-dependent; because ATR inhibition alone increases replication perturbation, however, we cannot determine whether ATR contributes to H2AX phosphorylation in the unperturbed subthreshold setting since its inhibition is known to stimulate ATM-dependent γH2AX (50). One possibility is that 100 nM APH preferentially affects regions prone to transcription–replication conflict, and another is that ATM senses an altered chromatin state rather than a lesion. Discriminating between these is a logical extension of this work but lies beyond its current scope.

The secretory output of the producer state is not a SASP. Indeed, induction is confined almost entirely to the protease and matrix-remodelling arm of the canonical SASP, while the NF-κB/IL-1/IL-6 core that defines it is not merely absent but co-ordinately repressed: TNFα signalling via NF-κB (NES - 1.70), IL6/JAK/STAT3 signalling (-1.75), IFN-γ (-1.61) and IFN-α (-1.55) responses are all significantly depleted. A conventional overlap test against a SASP gene list will therefore return a significant enrichment, driven entirely by the protease module. The producers, moreover, are demonstrably not senescent by SA-β-Gal, EdU, CDKN1A/CDKN2A expression and LMNB1. The comparison with genuine RS is equally informative: etoposide induces p53-p21, TNFα/ILs responses and represses proliferation genes, so the two transcriptional states are not graded versions of one another but move in opposite directions. Subthreshold perturbation of replication therefore defines an unexpected cell-autonomous response rather than a diluted RS/damage response.

Producer cells repress the matrisome as a whole and induce no TGF-β ligand, yet the responder cells are signalled to build matrix (matrisome NES +2.26; myCAF +2.44) and activate TGF-β signalling, upregulating both pathway components and latent TGF-β binding proteins. The bystander phenotype is therefore not a phenocopy of the producer phenotype, and TGF-β1 in this system is a recipient- induced output rather than the transmitted signal as it has been reported in other setting of genuine RS/DNA damage (51, 52). Similarly, although ATM and H2AX activation is mirrored in both compartments, as in RIBE, only the responder compartment converts that state into a matrix program, which implies that ATM activation, whatever its origin, is not sufficient on its own to specify the fibrogenic output.

The closest known relative of PeRBE is RIBE, and the consequences are inverted. RIBE is driven by DNA damage, involves ROS and gap-junctional contact, and delivers to bystander cells outcomes that are deleterious for them – DNA damage, cell-cycle arrest and eventually death (18, 19). PeRBE requires no DNA damage in the producer and is entirely ROS-independent. Where classical bystander effects propagate damage, PeRBE propagates an instruction; and the outcome in recipient cells is neutral or mildly pro-proliferative. Conversion of fibroblasts into ECM-building cells is generally referred to as fibroblast activation (26, 29), a state canonically induced by TGF-β, by mechanical stress or by inflammatory stimuli including those of the SASP (26, 53, 54). Our producer cells do not produce TGF-β, which cannot be the transmitted signal of PeRBE but responder cells clearly activate TGF-β signalling. ATM can license TGF-β under DNA damage or through production of ROS (55, 56). If in our responder cells ATM activates TGF-β, that would occur independent from ROS and DNA damage defining a distinct, physiological, rather than authentic pathological mechanism. Similarly, in cancer cells, ATM loss can stimulate myCAF (57) while our responder fibroblasts show up ATM phosphorylation and activate. From our data, we cannot conclude that PeRBE operates independently of TGF-β but we can support a model whereby PeRBE is a novel upstream trigger that converges on a fibrogenic program from a subthreshold perturbation of DNA replication, rather than injury, senescence, DNA damage, inflammation or mechanical stress, and characterized by non-senescent producers and non-arrested responders without genuine DNA damage and DDR. To our knowledge this is the first evidence linking the replication program itself to fibroblast activation *in trans* independently of DNA damage.

Chronic, low-grade perturbation of replication is expected in tissues experiencing oncogene activation in early lesions, hypomorphic variation in replication caretakers, metabolic imbalance or exposure to low doses of chemotherapeutics. If a perturbation of this magnitude is sufficient to instruct an ECM-producing state in neighbouring cells, replication dynamics may contribute to microenvironmental remodelling to establish a pro-fibrotic environment in the absence of a canonical, cellular pathology. Of note, the secretome of senescent cells is not a single stereotyped output being able to deliver different cellular changes. NOTCH1 activity switches oncogene-induced senescent cells between a TGF-β-rich and a pro-inflammatory secretome, with the two arms mutually antagonistic (58), and senescent cells at wound sites initiate myofibroblast differentiation through PDGF-AA rather than through the inflammatory core (59). On the other hand, senescent activated stellate cells secrete a protease-dominant, matrisome-repressed output that restrains rather than promotes fibrosis (60). Viewed against this heterogeneity, the PeRBE producer state resembles a single module – the protease and matrix-remodelling arm – deployed alone, in proliferating cells, with the inflammatory core not just unengaged but repressed. Whether this route contributes to fibrotic disease or to the fibroblast reprogramming that accompanies early tumorigenesis is an obvious question, which is outside the scope of this work.

This study has some limitations. The mediator or mediators of PeRBE are not identified beyond their biochemical properties, and the secretome is inferred largely from transcript data; how ATM-H2AX is activated in either compartment is not resolved, and we have not established whether responder ATM activation is required for the matrix programme; the phenomenon is characterised at a single dose and a single time point, in cultured fibroblasts; and the persistence, reversibility and possible onward propagation of the responder state remain untested and will deserve future studies to be addressed.

Nonetheless, the phenomenon we describe is distinct from both the SASP and RIBE, and reveals that a perturbation of DNA replication too mild to engage the checkpoint or to damage the cell in which it occurs generates a transmissible protein signal that converts naïve fibroblasts into proliferating matrix producers (Fig. 8F). That subtle shifts in replication dynamics can serve as tissue-level instructions reveals that the replication program functions as an intercellular signalling mechanism, expanding our understanding beyond cell-autonomous regulation.

## ACKNOWLEDGMENTS

Graphical abstract Created in BioRender. Pichierri, P. (2026) https://BioRender.com/2a5ae29 AUTHOR CONTRIBUTIONS: B.P. performed all the cell biology experiments for functional characterisation of the phenomenon. F.C. contributed to the characterization of the responder cell’s phenotype, proliferation and analysis of collagen production. F.D.F. performed the DNA fibres experiments. L.L.P. supervised the bioinformatics analyses and performed dataset comparisons. A.P. performed bioinformatics analysis. P.V. performed analysis of chromosomal damage. M.R. performed IncuCyte experiments. D.M supervised IncuCyte analysis. A.F. supervised chromosomal damage analysis and experiments with CFS-inducing APH dose. B.P., F.C., F.D.F., L.L.P., A.P., P.F., analysed data and contributed to designing the experiments and writing the paper. A.F., and P.P designed experiments, analysed data, supervised research and wrote the paper. All authors approved the paper.

## MATERIALS AND METHODS

### Cell lines

Primary human dermal fibroblasts (Ag21862) were used as the main cell line throughout the study; in selected experiments, a second primary fibroblast strain (Ag21859) was additionally used, as indicated in the corresponding figure legends. Cells were obtained from the Coriell Cell Repositories – NIA collection and maintained in Dulbecco’s Modified Eagle Medium (DMEM) supplemented with 10% fetal bovine serum (FBS) and cultured at 37 °C in a humidified atmosphere containing 5% CO₂ and 95% air. Cells were routinely passaged before reaching confluence and were regularly monitored and screened for mycoplasma contamination.

### Chemicals and reagents

The following compounds were used in this study:

- **Aphidicolin** from Sigma-Aldrich, cat. #178273; dissolved in DMSO;
- **Camptothecin** from Enzo Biochem, cat. #ALX-350-015; dissolved in DMSO;
- **Etoposide** from Selleck, cat. #S1225; dissolved in DMSO;
- **Colcemid** from Gibco KaryoMAX, cat. #15212012; supplied as a ready-to-use solution in PBS;
- **ATM Inhibitor** (KU-55933) from Selleck, cat. #S1092; dissolved in DMSO;
- **ATR Inhibitor** (VE-821) from Selleck, cat. #S8007; dissolved in DMSO;
- **DNAPKcs Inhibitor** (NU7441) from Selleck, cat. #S2638; dissolved in DMSO.

Stock solutions were diluted in complete culture medium to the final working concentrations indicated for each experiment.

### Experimental design and conditioned medium generation

To model the paracrine effects of a sub-threshold perturbation of DNA replication (PeRBE), a producer/responder co-culture-free system based on conditioned medium (CM) transfer was employed (Figure 6A). Producer fibroblasts were independently treated for 24 h with low, sub-cytotoxic doses of the DNA replication inhibitor aphidicolin (APH; 50, 100 or 400 nM), or with the topoisomerase inhibitors camptothecin (CPT; 50 nM) or etoposide (ETP; 50 µM), used as reference genotoxic/replication-stress-inducing agents; untreated producer cells (CTRL) were processed in parallel. Following treatment, cells were washed twice with PBS to remove residual drug and incubated in serum-free medium for an additional 24 h to allow the accumulation of secreted factors.

The conditioned medium was then collected, clarified, and diluted 1:1 with fresh serum-containing medium before being transferred onto untreated, naïve responder fibroblasts. Responder cells were exposed to the corresponding CM (CTRL CM, APH 50/100/400 nM CM, CPT 50 nM CM, or ETP 50 µM CM) for 24 h, unless otherwise specified, where responder cells were exposed to CM for 48 or 72 h to assess later time-point outcomes.

### Senescence-associated beta-galactosidase (SA β-GAL) activity assay

Cellular senescence was evaluated by measuring senescence-associated β-galactosidase (SA-β-gal) activity using the chromogenic substrate 5-bromo-4-chloro-3-indolyl β D-galactopyranoside (X-gal), as described (61). Briefly, cells were seeded in 24-well plates. After treatment, cells were washed twice with PBS and fixed with 4% paraformaldehyde for 5 min at room temperature. Following two washes with PBS, cells were incubated with SA-β-gal staining solution containing 0.1% X-gal, 5 mM potassium ferrocyanide, 5 mM potassium ferricyanide, 150 mM NaCl and 2 mM MgCl₂ in 40 mM citric acid/sodium phosphate buffer (pH 6.0). Cells were incubated at 37 °C in the dark until the development of the characteristic blue-green staining. The reaction was terminated by washing the cells with distilled water, and stained cells were visualized by bright-field microscopy using a 10× objective. SA-β-gal-positive cells were quantified as the percentage of blue-stained cells relative to the total number of cells analysed.

### EdU incorporation assay

The fraction of proliferating cells (% EdU-positive nuclei) was assessed by pulse labelling with the thymidine analogue EdU (15 µM), either for 30 min prior to harvesting or for 24 h concurrently with treatment, depending on the experiment. Detection was performed using the Click-iT EdU Imaging Kit (Invitrogen), according to the manufacturer’s specifications. Coverslips were observed at 20× with an Eclipse 80i Nikon fluorescence microscope equipped with a ViCo system. At least 200 nuclei were examined from three biological replicates.

### DNA Fiber Analysis

Replication fork progression was assessed using the DNA fiber assay. Following the indicated treatment, cells were pulse-labeled with 100 μM chlorodeoxyuridine (CldU), washed thoroughly with warm PBS and then incubated for an additional 30 min in fresh culture medium containing.25 μM iododeoxyuridine (IdU) for 30 min. After labeling, cells were harvested by trypsinization and cell pellets were resuspended in ice-cold PBS. A volume of 2–4 μL of the ice-cold cell suspension was spotted onto the upper end of a microscope slide and allowed to dry for 30 s. Subsequently, 12 μL of freshly prepared lysis buffer (200 mM Tris-HCl, pH 7.5, 50 mM EDTA, and 0.5% SDS) was added and gently mixed using a pipette tip. The slides were tilted at an angle of 25–40° for 2 min to allow the DNA to spread along the slide surface. Slides were then placed horizontally and allowed to air-dry completely in the dark for approximately 5–15 min.

Once dry, the slides were fixed in freshly prepared ethanol/acetic acid (3:1, v/v) for 15 min and subsequently allowed to dry. DNA replication tracks were detected by immunofluorescence using rat anti-CldU/BrdU (Abcam) and mouse anti-IdU/BrdU (BD Biosciences) primary antibodies, followed by appropriate fluorescently conjugated secondary antibodies. Images were acquired randomly from fields containing well-separated and non-overlapping DNA fibres using a Keyence BZ-X800 microscope. The lengths of the IdU- and CldU-labelled DNA tracks were measured using ImageJ software. Replication fork speed was calculated from the length of the labelled tracks and the corresponding labelling time, after conversion of track length from µm to kb using the appropriate DNA conversion factor.

For each experimental condition, at least 50 individual replication fibres/forks were analysed per experiment. Replication fork speed was quantified for each individual fork, and the distribution and percentage of forks displaying different replication speeds were determined for each experimental condition. In addition, replication fork symmetry was assessed by analysing pairs of sister forks originating from the same replication origin. The lengths of the two sister forks were measured independently, and forks were classified as symmetric or asymmetric according to the predefined difference between the two sister-fork progression rates. The percentage of symmetric and asymmetric replication forks was calculated for each experimental condition.

All measurements were performed using the same acquisition and analysis criteria across experimental groups. Each experiment was performed using three independent biological replicates.

### Western blotting analysis

Western blotting was performed using standard methods. The blots were incubated with the following primary antibodies: mouse anti-GAPDH (Millipore, 1:5000), mouse anti-p15 INK4B/p16 INK4A (C- 7) (Santa Cruz Biotechnology, 1:5000), rabbit anti-p21 Waf1/Cip1 (12D1) (Cell Signaling, 1:5000), mouse anti-CHK1 (Santa Cruz Biotechnology, 1:300), rabbit anti-p CHK1 (Cell Signaling, 1:1000), mouse anti- CHK2 (Cell Signaling, 1:1000), rabbit anti-p CHK2 (Cell Signaling, 1:1000), mouse anti-GAPDH (Millipore, 1:5000), mouse anti-COL1A (Santa Cruz Biotechnology, 1:1000).

After incubation with horseradish peroxidase-linked secondary antibodies (1:30 000, Jackson Immunoscience), the blots were detected using the Western Bright ECL detection kit (Advansta) according to the manufacturer’s instructions. Quantification was performed on scanned images of the blots using Image Lab software, with values shown on the graphs normalized to protein content as evaluated through GAPDH.

### Detection of ssDNA by native IdU assay

To detect parental ssDNA, cells were labelled for 20 hours with 80 μM IdU (Sigma-Aldrich), released in fresh DMEM for 2 hours, then treated as indicated. For immunofluorescence, cells were washed with PBS 1X, permeabilized with 0.5% Triton X-100 for 10 min at 4 °C and fixed in 3% PFA, 2% sucrose in PBS 1X. Fixed cells were then incubated with mouse anti-IdU antibody (Becton Dickinson, 1:80) for 1 h at 37 °C in 1% BSA/PBS, followed by species-specific fluorescein-conjugated secondary antibodies (Alexa Fluor 488 Goat Anti-Mouse IgG (H + L), highly cross-adsorbed—Life Technologies). Slides were analysed with Eclipse 80i Nikon Fluorescence Microscope, equipped with a Virtual Confocal (ViCo) system. For each time point, at least 100 nuclei were analysed. Quantification was carried out using the ImageJ software.

### Immunofluorescence assays

Cells were grown on 35-mm coverslips and harvested at the indicated times after treatments. For γH2AX, p-ATM, and p-KAP1 IF, cells were washed with PBS, pre-extracted with 0.5% Triton X- 100, and fixed with 4% PFA at RT for 10 min. After blocking in 3% BSA for 20 min, staining was performed with mouse anti-γH2AX (S139) (Millipore, 1:1000), rabbit anti-phospho-KAP1 (S824) (Bethyl, 1:500), or mouse anti-phospho-ATM (Millipore, 1:300), diluted in 1% BSA/0.1% saponin in PBS, for 1 h at room temperature in a humidified chamber.

For COL1A immunofluorescence, cells were fixed with 4% PFA at +4 °C for 10 min, permeabilized with 0.1% Triton-X in PBS for 10 min at RT and blocked in 1% BSA for 1 h at RT. The primary antibody anti-COL1A (Santa Cruz Biotechnology) was diluted 1:200 in 0.1% BSA in PBS and incubated ON at + 4°C in a humidified chamber,

After each primary antibody incubation, cells were washed twice with PBS and incubated with the appropriate secondary antibody — goat anti-mouse Alexa Fluor 488 or goat anti-rabbit Alexa Fluor 594 (Molecular Probes), both diluted 1:200 — for 1 h at RT in a humidified chamber. For COL1A immunofluorescence, cells were incubated with goat anti-mouse Alexa Fluor 488 diluted 1:500 in 0.1% BSA in PBS and incubated for 1 h at RT in a humidified chamber. DNA was counterstained with 0.5 μg/mL DAPI. Images were randomly acquired using an Eclipse 80i Nikon fluorescence microscope equipped with a ViCo system. For each condition, at least 200 nuclei were acquired at 40× magnification. Only nuclei with >5 foci were considered positive and were quantified using ImageJ. Quantification of COL1A fluorescence signal was performed using ImageJ software following the protocol for Corrected Total Cell Fluorescence (CTCF) (*Measuring cell fluorescence using ImageJ.* The Open Lab Book v1.0. Available online: https://theolb.readthedocs.io/en/latest/imaging/measuring-cell-fluorescence-using-imagej.html). CTCF was calculated using the formula: CTCF = Integrated Density - (Cell Area x Mean Background Fluorescence). Data were expressed as COL1A fluorescence intensity in arbitrary units (A.U.).

### Comet assay

DNA damage was assessed by alkaline and neutral Comet assay (single-cell gel electrophoresis) (34, 62). In both cases, cells were embedded in low-melting-point agarose and spread onto glass slides; for the neutral assay, slides were subjected to electrophoresis (20 min, 6–7 A, 20 V) and fixed in methanol. DNA was stained with GelRed (Biotium) and comets were visualized using an Olympus fluorescence microscope (20× magnification for the alkaline assay, 20× for the neutral assay). Slides were analysed with a computerized image analysis system (CometScore, Tritek Corp.). The extent of DNA damage was quantified as tail moment (tail length × fraction of total DNA in the tail). Apoptotic cells, identified by a small comet head and disproportionately large tail, were excluded from the analysis to avoid artificial inflation of tail moment values.

### In situ proximity-ligation assay (PLA)

Cells were cultured on 8-well chamber slides (Millicell, Sigma-Aldrich). After the indicated treatments for 24 h, cells were pre-extracted with 0.5% Triton X-100 for 8 min on ice, fixed with 3% PFA/2% sucrose in 1X PBS for 15 min at room temperature (RT), permeabilized with 0.25% Triton X-100 for 15 min at RT, and blocked in blocking buffer. The in-situ proximity ligation assay (PLA) was performed using the NaveniFlex kit (Navinci Diagnostics) with anti-Mouse PLUS and anti-Rabbit MINUS PLA probes, according to the manufacturer’s instructions. To detect the proteins of interest, mouse anti-phospho-ATM (Millipore, 1:600) and rabbit anti-ATM (Abclonal, 1:400) antibodies were used. Ligation and amplification were performed at 37°C. PLA foci were visualized by fluorescence microscopy and quantified using ImageJ.

### Chromosomal aberrations

Chromosomal aberrations were induced by treating cells with 50, 100, or 400 nM aphidicolin (APH) for 24 hours. Cell cultures were incubated with 0.1 µg/mL Colcemid at 37°C for 3–6 h until harvesting. Cells for metaphase preparations were collected according to standard procedure (34). In brief, the cellular pellet was resuspended in prewarmed hypotonic solution (0.075 M KCl in distilled water) and incubated at 37°C for 20 min, followed by multiple changes of fixative solution (3:1 methanol/acetic acid). The cell suspension was dropped onto cold, wet slides to prepare chromosome spreads. Slides were air dried overnight and stored at −20°C until analysis. For each treatment condition, the number of breaks and gaps was scored on Giemsa-stained metaphases. For each time point, a minimum of 50 metaphases were independently examined by two investigators, and chromosomal damage was scored at 100× magnification using an Olympus fluorescence microscope.

### Live-imaging analysis

Cells were seeded on 24-well plates and treated as described above. At time zero (T0) plates were placed into the Incucyte® SX5 in a humified CO2 incubator at 36.5°C. Scanning was scheduled every 4 hours for 5 days with the brightfield filter at 10× magnification. At the end of the acquisition, a sample of positive (fully confluent at day 5) and negative (low confluence at T0) control image were used to setup the analysis pipeline by using the Incucyte® SX5 Analysis Software. Confluence area were calculated and, to avoid artifacts due to seeding variability, it was normalized versus the T0 for each sample and expressed as area units. Growth rate and flex point were calculated using GraphPad Prism Software through non-linear regression analysis and curve fitting to a logistic growth equation.

### RNA extraction

Total RNA was extracted from fibroblasts (cell line 21862) using the Quick-RNA™ Miniprep Kit (Zymo Research, Irvine, CA, USA; cat. R1054), according to the manufacturer’s instructions. Cells were lysed directly using the provided lysis buffer, and RNA was subsequently purified by adsorption onto silica matrix spin columns (Zymo-Spin), exploiting the kit’s proprietary technology that enables rapid isolation of total RNA free of genomic DNA contamination, aided by an on-column DNase I treatment. RNA concentration was determined by fluorometry (Qubit™ RNA Assay Kit, Thermo Fisher Scientific), while RNA integrity was assessed by automated capillary electrophoresis (Agilent High Sensitivity RNA ScreenTape, TapeStation), calculating the RNA Integrity Number (RIN). All samples showed a high degree of integrity, with RIN values ranging from 9.0 to 9.7, indicating excellent RNA quality suitable for downstream analyses. RNA samples were stored at −80 °C until use.

### RNA quality control, library preparation, and sequencing

Total RNA extraction was performed on four biological replicates per condition: A) baseline cells, B) cells treated with 100 nM Aphidicolin, C) cells conditioned with medium from baseline cells, D) cells conditioned with medium from treated cells. Sample quality control, library preparation, and high-throughput sequencing were performed by Novogene Co., Ltd. (Beijing, China). RNA integrity, purity, and concentration were evaluated prior to library construction using a NanoDrop 2000 spectrophotometer and an Agilent Bioanalyzer 2100 system. Only high-quality RNA samples with an RNA Integrity Number (RIN) > 7.5 were subjected to library construction. Sequencing libraries were constructed following standard Illumina protocols (NEBNext Ultra RNA Library Prep Kit). The resulting libraries were multiplexed and sequenced on an Illumina NovaSeq X Plus platform to generate 150 bp paired-end (PE150) reads, targeting a depth range of 80-91 million reads per sample.

### Data preprocessing, alignment, and quantification

Raw sequencing data in FASTQ format were preprocessed to remove adapter sequences and low-quality reads (bases with a Phred quality score Q<=20 accounting for >50% of the read). High-quality clean paired-end reads were mapped to the human reference genome GRCh38 (Ensembl primary assembly (63) using HISAT2 (v2.2.1) (64). Transcript assembly and gene-level expression quantification were computed using StringTie (v2.2.3) (65) in reference-only mode guided by Ensembl GRCh38 gene annotation (Ensembl release 102) (63). The resulting raw read count matrix was exported for downstream analysis.

### Differential gene expression analysis

Statistical analysis was performed in the R software environment (v4.4.1) using the edgeR package (v4.4.2) [4]. Low-abundance genes were filtered out by retaining only those with counts per million (CPM) >1 in at least 4 samples. Library size normalization was applied using the Trimmed Mean of M-values (TMM) method. Pairwise differential gene expression between experimental conditions was evaluated using the quasi-likelihood F-test implemented in edgeR. P-values were adjusted for multiple testing using the Benjamini-Hochberg False Discovery Rate (FDR) procedure. Genes exhibiting an adjusted P-value (FDR) < 0.05 and an absolute magnitude of |log_2_(FoldChange)| >= 1.0 were considered significantly differentially expressed genes (DEGs).

### Sensitivity analyses

For the producer dataset, a replicate-blocked model (∼replicate + condition) was fitted as a sensitivity analysis and returned an essentially identical gene set (1957 versus 2014 differentially expressed genes), indicating no replicate pairing structure. For the responder dataset, in which the control group showed greater heterogeneity, models were refitted after removing the one and the two most divergent control replicates; these returned 1063 and 564 differentially expressed genes with gene-level log2 fold changes correlating with the full model at r = 0.965 and r = 0.835 respectively. Effect directions were preserved for every gene class in every model, whereas effect magnitudes were reduced by approximately half in the most conservative model.

### Functional Enrichment Analysis

Analyses used MSigDB v7.0 (Hallmark, C2 canonical pathways including Reactome, C5 Gene Ontology Biological Process, and C3 transcription-factor-target collections). The following additional sets were defined:

The matrisome was defined as NABA_MATRISOME. The secreted and extracellular compartment was defined as the union of the matrisome with GO_CYTOKINE_ACTIVITY, GO_CHEMOKINE_ACTIVITY, GO_GROWTH_FACTOR_ACTIVITY, GO_EXTRACELLULAR_MATRIX and GO_COLLAGEN_CONTAINING_EXTRACELLULAR_MATRIX. A core senescence-associated secretory phenotype set was curated from the literature and organised into six functional modules (IL- 1/IL-6 core inflammatory, chemokines, IGF axis, proteases and inhibitors, growth factors and other soluble mediators, and innate immune and acute-phase factors); of its members, 75 were detected in the producer dataset. For cross-checking, a broader SASP definition was used, comprising the union of this curated set with REACTOME_SENESCENCE_ASSOCIATED_SECRETORY_PHENOTYPE_SASP and FRIDMAN_SENESCENCE_UP. Additional curated sets comprised canonical p53 target genes, chromatin writers and histone chaperones, a myofibroblast/fibrotic ECM (myCAF) signature, cytoplasmic ribosomal protein genes, replication-dependent histone genes and replication-independent histone variants, the latter two curated separately so that the S-phase specificity of the histone response could be tested. A 198-gene replication-stress and genome-maintenance panel was assembled in ten modules (Fanconi anaemia pathway; homologous recombination and fork protection; RecQ helicases and fork remodelling; structure-specific nucleases and resolution; ATR- CHK1 checkpoint; translesion synthesis and fork restart; replisome and origin licensing; dNTP supply; R-loop and RNA:DNA hybrid control; interstrand crosslink and end processing). All curated sets is provided in the supplementary tables.

Functional enrichment analysis across multiple biological databases (including Gene Ontology (GO) (40) categories (Biological Process, Molecular Function, Cellular Component), KEGG Reactome (66) and WikiPathways (67), was performed using g:Profiler (68) (g:GOSt functional profiling tool) and R-based enrichment visualization packages. Input lists of DEGs were mapped to their corresponding Ensembl Gene IDs (ENSG). Non-coding transcripts (such as antisense RNAs and lncRNAs) and unannotated genes were automatically filtered out by g:Profiler based on the selected target organism database and functional annotations. Significance threshold for enriched terms and pathways was defined using Benjamini-Hochberg FDR < 0.05.

### Over-representation analysis

Over-representation of gene sets among the up- and down-regulated gene lists was assessed by the hypergeometric test, using all tested genes as the background universe and requiring at least three overlapping genes and a gene set size between 10 and 1000. P values were adjusted by the Benjamini– Hochberg procedure. Enrichment of differentially expressed genes within defined gene classes was tested by two-sided Fisher’s exact test against the background rate for the same dataset, and class-wise distributions of effect sizes were compared with the two-sided Mann–Whitney U test against all tested genes.

### Signature scoring

Per-replicate signature scores were computed by converting variance-stabilised counts to gene-wise z-scores across all samples of the relevant experiment and averaging these across the members of each gene set detected in that dataset. Scores were compared between conditions with the two-sided Mann–Whitney U test. With four replicates per group the minimum attainable P value for this test is 0.029; where this value is reported, the substantive observation is the complete separation of replicates between groups.

### Etoposide dataset

The etoposide comparison (50 nM, 24 h, versus untreated) was supplied as a processed differential expression table containing effect sizes, nominal and adjusted P values, an F statistic and per-group normalised counts per million. In the file as supplied, the reported log2 fold change is oriented as untreated over etoposide; effect sizes were therefore inverted before analysis so that positive values denote induction by etoposide, and the orientation was confirmed independently from the per-group normalised counts and from the direction of canonical p53 target and proliferation genes. All comparisons with the aphidicolin datasets were performed on gene symbols, restricted to genes tested in both experiments, and using the significance thresholds defined above.

### Statistical analysis

Experiments shown are representative of at least three independent biological replicates. Statistical differences were determined using the built-in tools in Prism 10 (GraphPad Inc.) by one- or two-way ANOVA, Mann-Whitney U-test or Student’s t-test. P < 0.05 was considered as significant.

