## Supplementary Figures and Legends for "Subthreshold perturbation of DNA replication induces a secretory response and a bystander effect in naïve human fibroblasts"

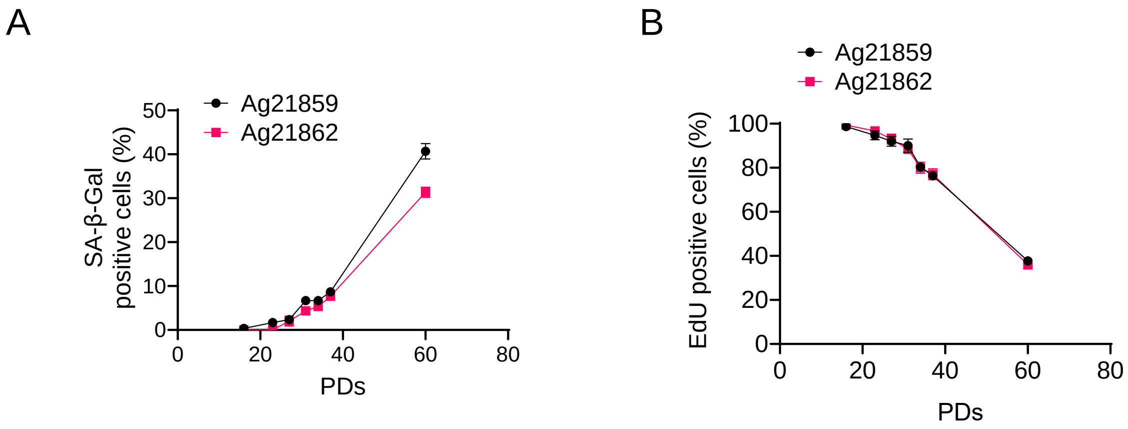
A
Ag21859
Ag21862
SA-β-Gal positive cells (%)
50
40
30
20
10
0
0
20
40
60
80
PDs
B
Ag21859
Ag21862
EdU positive cells (%)
100
80
60
40
20
0
0
20
40
60
80
PDs

Figure S1

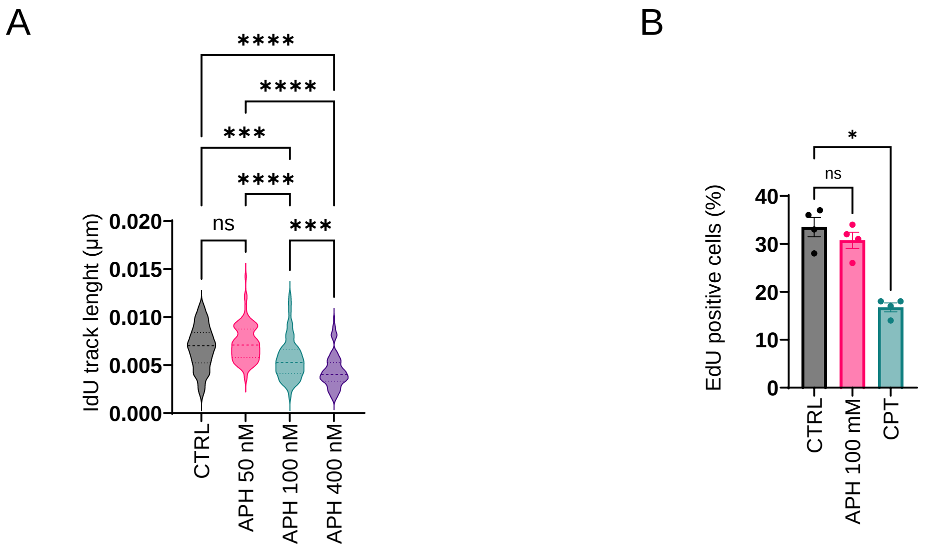
A
****
****
***
****
ns
***
IdU track lenght (µm)
0.020
0.015
0.010
0.005
0.000
CTRL
APH 50 nM
APH 100 nM
APH 400 nM
B
*
ns
EdU positive cells (%)
40
30
20
10
0
CTRL
APH 100 mM
CPT

Figure S2

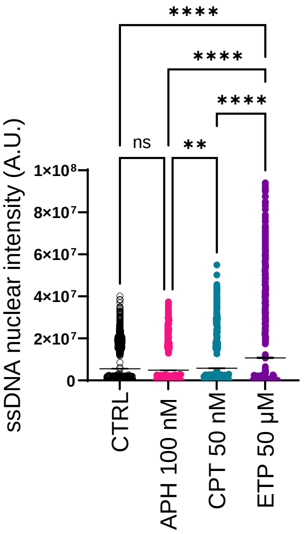
****
****
****
ns
**
ssDNA nuclear intensity (A.U.)
1×10⁸
8×10⁷
6×10⁷
4×10⁷
2×10⁷
0
CTRL
APH 100 nM
CPT 50 nM
ETP 50 μM

Figure S3

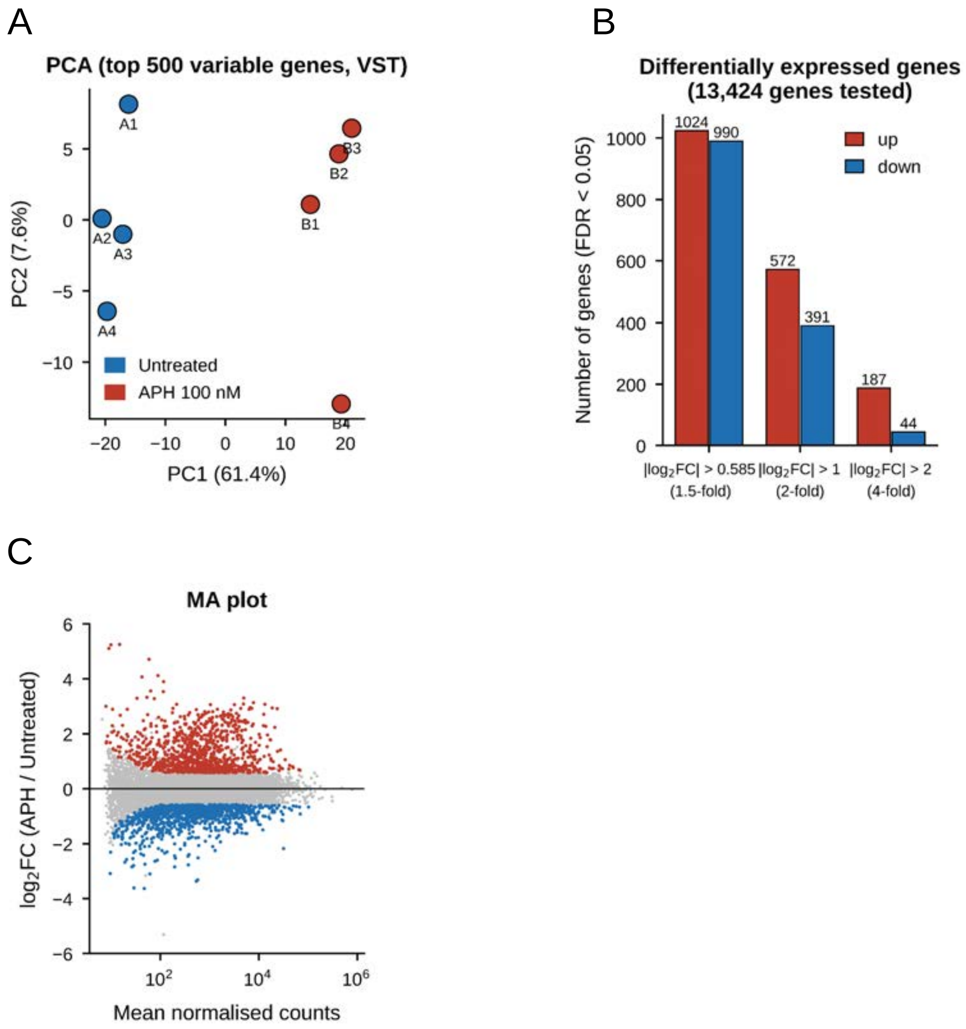
A
PCA (top 500 variable genes, VST)
PC2 (7.6%)
PC1 (61.4%)
Untreated
APH 100 nM
A1
A2
A3
A4
B1
B2
B3
B4
B
Differentially expressed genes
(13,424 genes tested)
Number of genes (FDR < 0.05)
up
down
1024
990
572
391
187
44
|log2FC| > 0.585
(1.5-fold)
|log2FC| > 1
(2-fold)
|log2FC| > 2
(4-fold)
C
MA plot
log2FC (APH / Untreated)
Mean normalised counts

Figure S4

Upregulated – APH vs. CTRL

A

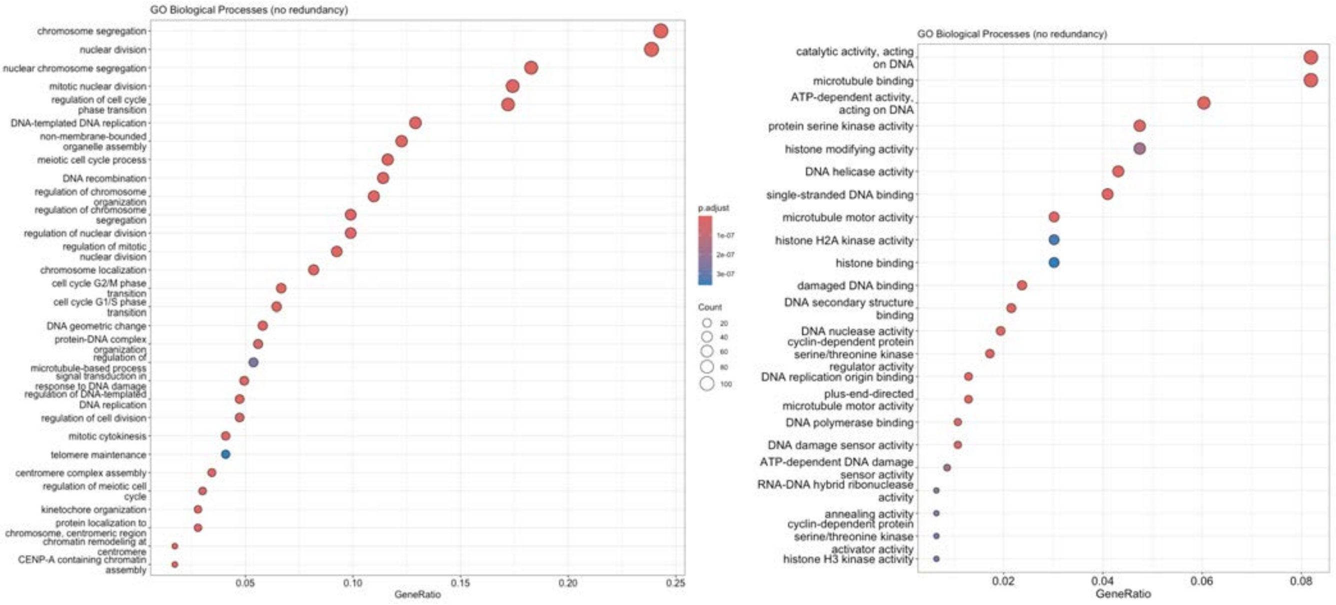


Downregulated – APH vs. CTRL

B

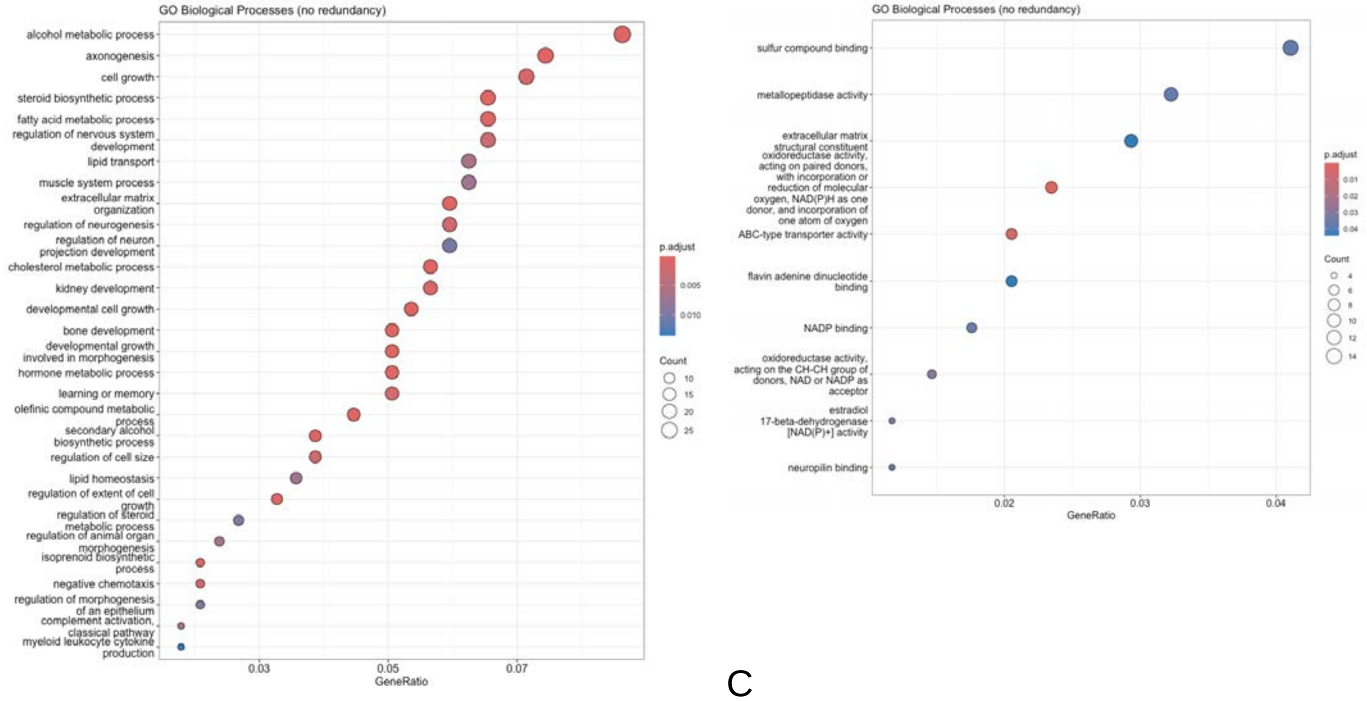


C

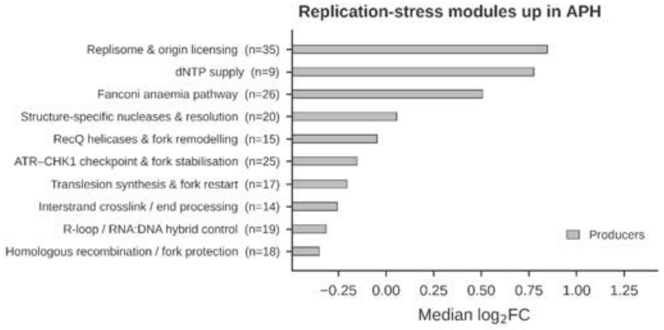


Figure S5

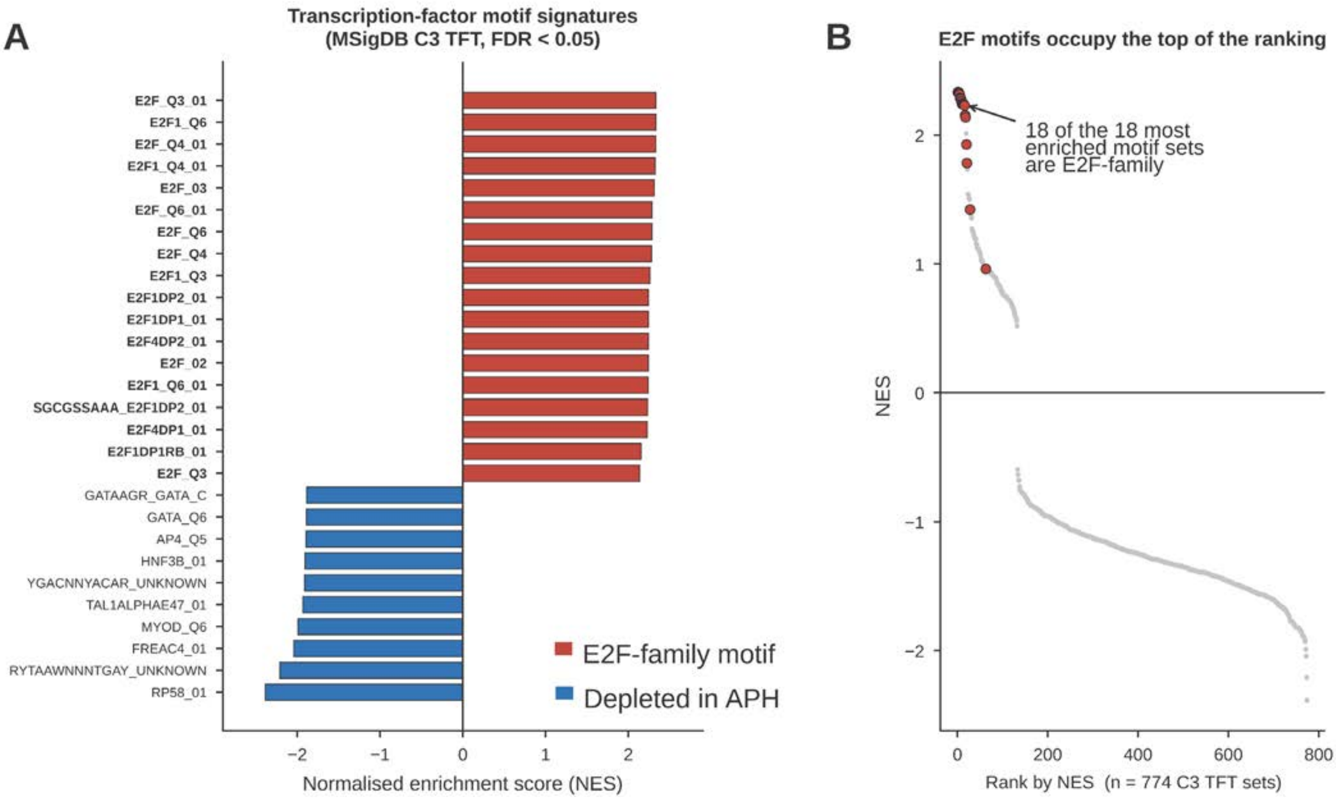
A
Transcription-factor motif signatures
(MSigDB C3 TFT, FDR < 0.05)
E2F_Q3_01
E2F1_Q6
E2F_Q4_01
E2F1_Q4_01
E2F_03
E2F_Q6_01
E2F_Q6
E2F_Q4
E2F1_Q3
E2F1DP2_01
E2F1DP1_01
E2F4DP2_01
E2F_02
E2F1_Q6_01
SGCGSSAAA_E2F1DP2_01
E2F4DP1_01
E2F1DP1RB_01
E2F_Q3
GATAAGR_GATA_C
GATA_Q6
AP4_Q5
HNF3B_01
YGACNNYACAR_UNKNOWN
TAL1ALPHAE47_01
MYOD_Q6
FREAC4_01
RYTAAWNNNTGAY_UNKNOWN
RP58_01
E2F-family motif
Depleted in APH
−2
−1
0
1
2
Normalised enrichment score (NES)
B
E2F motifs occupy the top of the ranking
18 of the 18 most enriched motif sets are E2F-family
NES
2
1
0
−1
−2
0
200
400
600
800
Rank by NES (n = 774 C3 TFT sets)

Figure S6

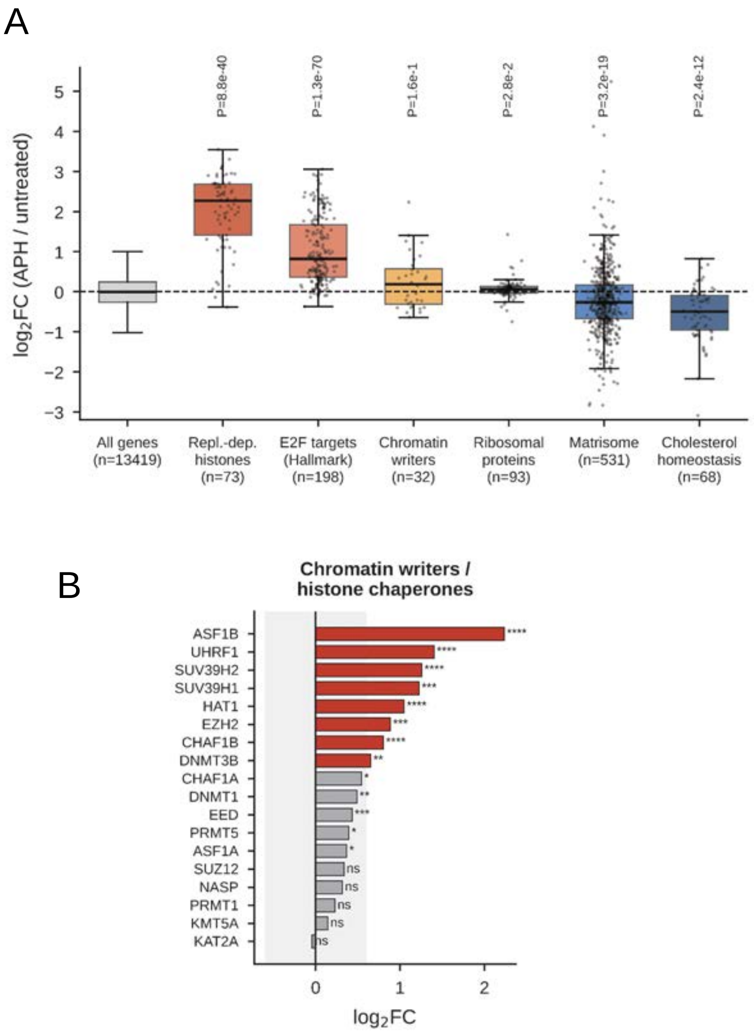
A
log2FC (APH / untreated)
P=8.8e-40
P=1.3e-70
P=1.6e-1
P=2.8e-2
P=3.2e-19
P=2.4e-12
All genes (n=13419)
Repl.-dep. histones (n=73)
E2F targets (Hallmark) (n=198)
Chromatin writers (n=32)
Ribosomal proteins (n=93)
Matrisome (n=531)
Cholesterol homeostasis (n=68)
B
Chromatin writers / histone chaperones
ASF1B ****
UHRF1 ****
SUV39H2 ****
SUV39H1 ***
HAT1 ****
EZH2 ***
CHAF1B ****
DNMT3B **
CHAF1A *
DNMT1 **
EED ***
PRMT5 *
ASF1A *
SUZ12 ns
NASP ns
PRMT1 ns
KMT5A ns
KAT2A ns
log2FC

Figure S7

A

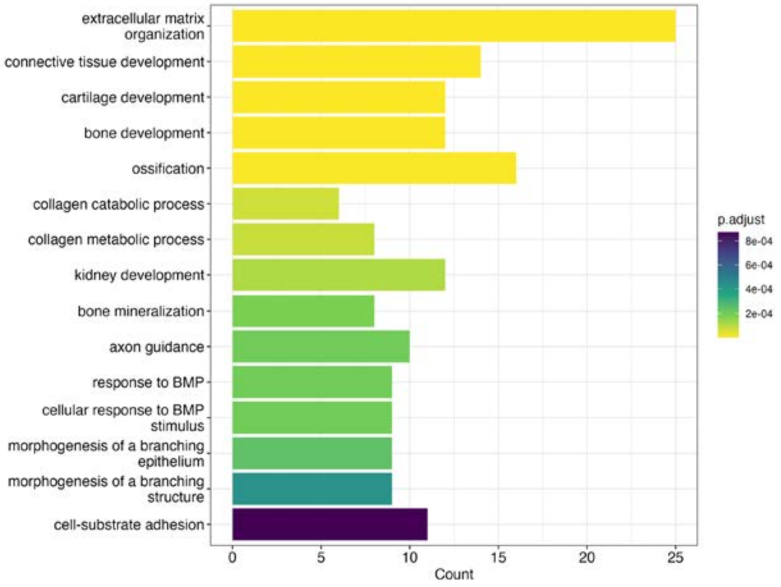
extracellular matrix organization
connective tissue development
cartilage development
bone development
ossification
collagen catabolic process
collagen metabolic process
kidney development
bone mineralization
axon guidance
response to BMP
cellular response to BMP stimulus
morphogenesis of a branching epithelium
morphogenesis of a branching structure
cell-substrate adhesion
0
5
10
15
20
25
Count
p.adjust
8e-04
6e-04
4e-04
2e-04

B

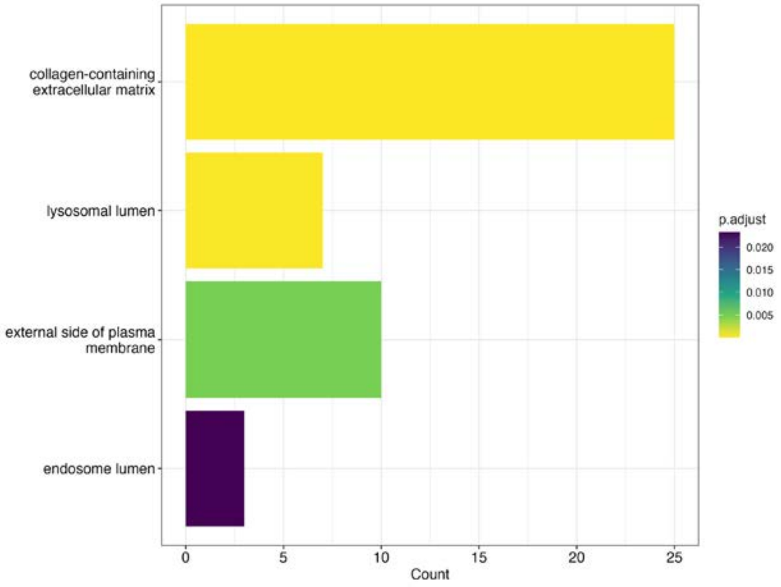
collagen-containing extracellular matrix
lysosomal lumen
external side of plasma membrane
endosome lumen
0
5
10
15
20
25
Count
p.adjust
0.020
0.015
0.010
0.005

Figure S8

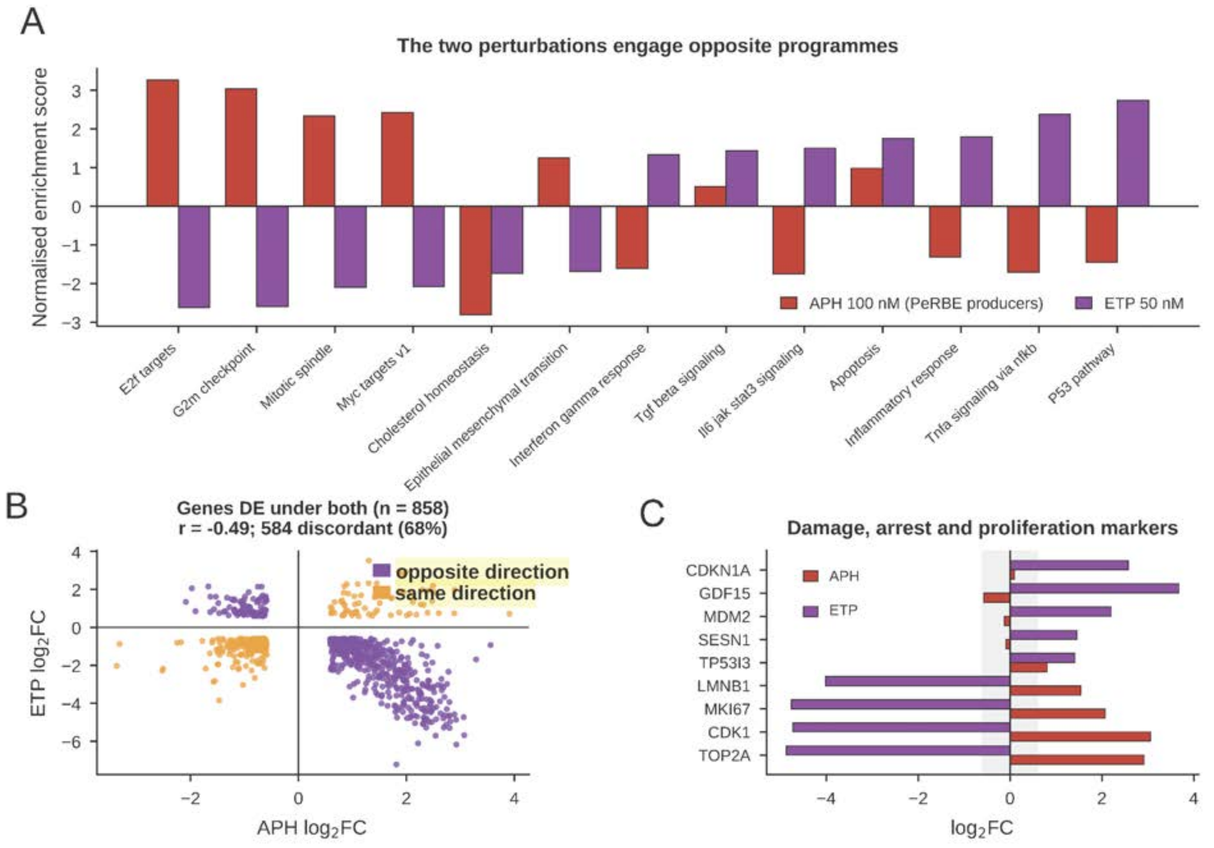
A
The two perturbations engage opposite programmes
Normalised enrichment score
APH 100 nM (PeRBE producers)
ETP 50 nM
E2f targets
G2m checkpoint
Mitotic spindle
Myc targets v1
Cholesterol homeostasis
Epithelial mesenchymal transition
Interferon gamma response
Tgf beta signaling
Il6 jak stat3 signaling
Apoptosis
Inflammatory response
Tnfa signaling via nfkb
P53 pathway
B
Genes DE under both (n = 858)
r = -0.49; 584 discordant (68%)
opposite direction
same direction
ETP log2FC
APH log2FC
C
Damage, arrest and proliferation markers
APH
ETP
CDKN1A
GDF15
MDM2
SESN1
TP53I3
LMNB1
MKI67
CDK1
TOP2A
log2FC

Figure S9

A

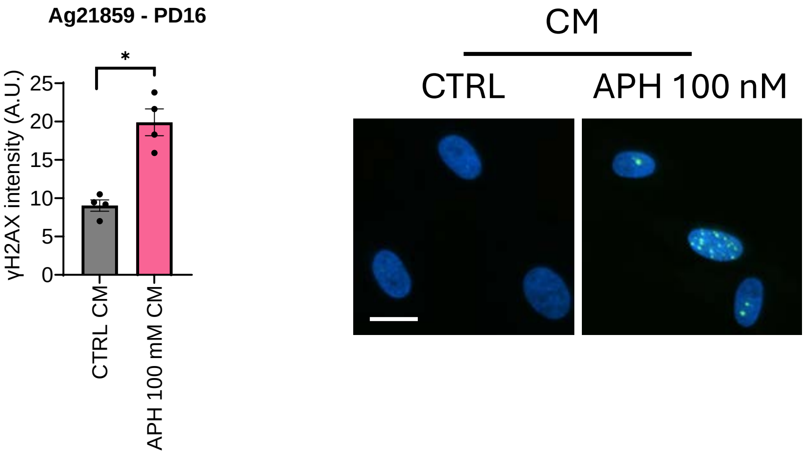


B

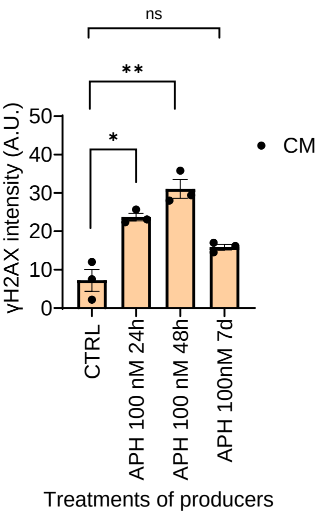


Figure S10

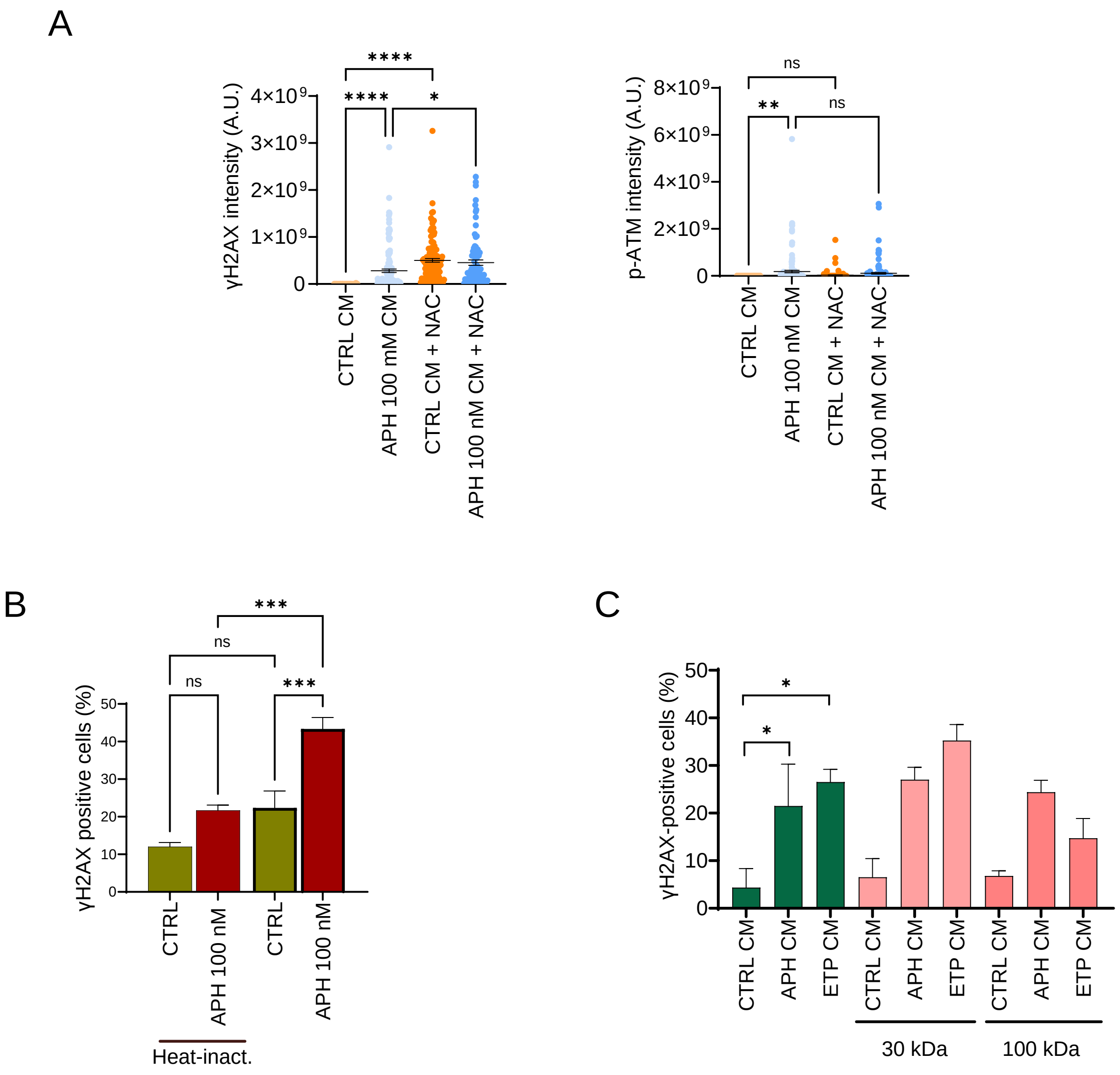

A
γH2AX intensity (A.U.)
0
1×10⁹
2×10⁹
3×10⁹
4×10⁹
****
****
*
CTRL CM
APH 100 mM CM
CTRL CM + NAC
APH 100 nM CM + NAC
p-ATM intensity (A.U.)
0
2×10⁹
4×10⁹
6×10⁹
8×10⁹
ns
**
ns
CTRL CM
APH 100 nM CM
CTRL CM + NAC
APH 100 nM CM + NAC
B
γH2AX positive cells (%)
0
10
20
30
40
50
***
ns
ns
***
CTRL
APH 100 nM
CTRL
APH 100 nM
Heat-inact.
C
γH2AX-positive cells (%)
0
10
20
30
40
50
*
*
CTRL CM
APH CM
ETP CM
CTRL CM
APH CM
ETP CM
CTRL CM
APH CM
ETP CM
30 kDa
100 kDa


Figure S11

SUPPLEMENTARY FIGURES LEGENDS

**Figure S1. Analysis of the proliferative potential of the fibroblast strains used in the study.**

The two healthy primary human fibroblasts were analysed at increasing population doublings (PD 16–60) for their proliferative potential. **(A)** Senescence, evaluated as the percentage of SA-β-galactosidase-positive cells. Data are expressed as mean ± SD (n = 3 biological replicates per strain). **(B)** Replicative potential, evaluated as the percentage of EdU-positive nuclei after a 24h labelling period. Data are expressed as mean ± SD (n = 3 biological replicates per strain).

**Figure S2. Subthreshold APH minimally affects fork progression and does not impair overall replication.**

**(A)** IdU track length (µm) in cells treated with 50, 100, or 400 nM APH for 24 h, or untreated. Statistical significance was determined by one-way ANOVA; ns, not significant ($p > 0.05$), *** $p < 0.001$, **** $p < 0.0001$. **(B)** EdU-positive cells (%) at the end of treatment with 100 nM APH or 50 nM CPT, compared to untreated cells. Statistical significance was determined by one-way ANOVA; ns, not significant ($p > 0.05$), * $p < 0.05$.

**Figure S3. Subthreshold APH does not induce parental ssDNA exposure.** Quantification of ssDNA nuclear intensity (A.U.) in cells treated with 100 nM APH, 50 nM CPT, or 50 µM ETP for 24 h, compared to untreated controls. Statistical significance was determined by one-way ANOVA; ns, not significant ($p > 0.05$), * $p < 0.05$, ** $p < 0.01$, **** $p < 0.0001$.

**Figure S4. Quality control of the producer cells RNAseq.**

**(A)** Principal component analysis (PCA) of variance-stabilised counts of the 500 most variable genes. **(B)** Pearson correlation between all pairs of samples at increasing values of significance and fold-increase. **(C)** MA plot of Log2 fold-change against mean normalised counts.

**Figure S5. Functional Gene Ontology (GO) enrichment analysis of the responder cells DEGs.**

**(A)** Dot plots representing biological processes (left panel) and molecular function categories (right panel) upregulated or downregulated **(B)** in APH-treated cells vs. CTRL. In BP dot plot, the horizontal axis represents the Gene Ratio (the proportion of differential genes mapped to the given pathway). Circle size corresponds to the number of genes assigned to each term (Count), while the colour gradient indicates statistical significance based on Benjamini-Hochberg adjusted p-values (p.adjust). In MF dot plot, terms are ordered by Gene Ratio along the horizontal axis. Dot sizes are proportional to gene counts involved in each functional category (Count), and dot colours reflect the significance levels according to adjusted *p*-values (p.adjust). **(C)** Bar chart showing replication stress-related genes modulated by subthreshold APH in producer cells shown as median log2 fold-change. The number of significantly-induced genes per module is indicated, including the Fanconi anaemia pathway, dNTP supply and metabolism, structure-specific nucleases and fork resolution, forkremodelling helicases, ATR-CHK1 signalling and activation, translesion synthesis and fork repair, R-loop resolution/fork protection, and homologous recombination/fork protection factors.

**Figure S6. Transcription-factor motif enrichment in producer cells is dominated by E2F-family motifs.**

**(A)** Normalised enrichment scores for the 18 most enriched and 10 most depleted MSigDB C3 transcription factor-target gene sets (Hallmark, Reactome, GO Biological Process, C3 transcription-factor targets) (FDR<0.05). E2F family members are shown in red/bold. **(B)** All 774 C3 TFT sets in APH-treated cells ranked by normalised enrichment score (E2F family highlighted). All 18 top-ranked sets are E2F-family E2F motifs account for 20 of the 24 significantly enriched sets.

**Figure S7. Subthreshold perturbation of DNA replication by APH induces a coordinated replication-coupled histone and chromatin-writer programs.**

**(A)** Distribution of log2 fold-changes by gene classes. Boxes show the median and interquartile range, whiskers extend to 1.5x the interquartile range, and dots are individual genes (P-values are from Mann-Whitney two-sided U-test against all genes). **(B)** Log2 fold-changes of chromatin writers and histone chaperones (ASF1B, UHRF1, SUV39H2, SUV39H1, HAT1, EZH2, CHAF1B, DNMT3B, CHAF1A, DNMT3A, EED, PRMT5, ASF1A, SUZ12, NASP, PRMT1, KMT5A, KAT2A). (Statistical significance is from Mann-Whitney two-sided U-test; ns, not significant ($p > 0.05$), * $p < 0.05$, ** $p < 0.01$, *** $p < 0.001$, **** $p < 0.0001$).

**Figure S8. Functional Gene Ontology (GO) enrichment analysis of the secretome-associated DEGs.**

Barplots representing top-15 enriched terms across the three GO categories for the 118 secretome candidate DEGs in producer cells (APH vs CTRL, filtered for the GO term GO:0005576 "extracellular region" and HPA secretome annotations): **(A)** Biological Process (BP) and **(B)** Cellular Component (CC). Bar lengths indicate the number of genes assigned to each term (Count), and colour gradient represents statistical significance based on Benjamini-Hochberg adjusted p-values (p.adjust).

**Figure S9. Genuine DNA damage/RS induces a p53/NF-κB inflammatory response that is transcriptionally opposite to the PeRBE producer state.**

**(A)** Paired Hallmark normalised enrichment scores for producer fibroblasts treated with 100 nM APH or with 50 nM ETP for 24 h. **(B)** Genes identified as differentially expressed under both treatments ($n = 858$); purple, opposite direction; amber, same direction; 68% change direction (Pearson $r = -0.50$). **(C)** Log2 fold-changes of representative DNA damage, cell-cycle arrest, and proliferation marker genes (CDKN1A, GDF15, MDM2, SESN1, TP53I3, LMNB1, MKI67, CDK1, TOP2A) in APH- (red) versus ETP-treated (purple) cells. Raw data are from Supplementary Table 2 (N=3).

**Figure S10. Bystander H2AX phosphorylation is reproducible in an independent fibroblast strain and depends on the duration of producer treatment.**

**(A)** Quantification (left) and representative immunofluorescence images (right) of γH2AX intensity in a second strain of young primary human fibroblasts (Ag21859, PD16) exposed to CM from control or APH-treated (100 nM) producer cells. Representative images are shown (Scale bar 20µm). **(B)** γH2AX intensity in responder cells exposed to CM collected from producer cells treated with 100 nM APH for 24 h, 48 h, or 7 days, showing that the bystander response declines with prolonged APH treatment of producers. Data are mean ± SD; ns, not significant; *P < 0.05; **P < 0.01.

**Figure S11. The PeRBE-transmitted signal is ROS-independent, proteinaceous, and in a defined range of molecular weight.**

**(A)** Quantification of γH2AX and p-ATM intensity in responder cells exposed to CM from control or APH-treated (100 nM) producers, in the presence or absence of the ROS scavenger NAC, showing that NAC treatment does not abrogate the bystander response. **(B)** Percentage of γH2AX-positive responder cells exposed to control or APH (100 nM) CM, either untreated or after heat inactivation of the CM (65°C), showing that heat inactivation substantially reduces bystander γH2AX induction. **(C)** Percentage of γH2AX-positive responder cells exposed to unfractionated CM or to CM size-fractionated through 30 kDa and 100 kDa cut-off filters, from control, APH-, or ETP-treated producers, showing that bystander activity is retained in both fractions and is enriched in the ≤30 kDa fraction. Data are mean ± SD; ns, not significant; *P < 0.05; **P < 0.01; ***P < 0.001; ****P < 0.0001.
